# Leave no stone unturned: The hidden potential of carbon and nitrogen cycling by novel, highly adapted *Thaumarchaeota* in the Atacama Desert hyperarid core

**DOI:** 10.1101/2020.07.17.208546

**Authors:** Yunha Hwang, Dirk Schulze-Makuch, Felix L. Arens, Johan S. Saenz, Panagiotis S. Adam, Till L. V. Bornemann, Alessandro Airo, Michael Schloter, Alexander J. Probst

## Abstract

The hyperarid core of the Atacama Desert is an extremely harsh environment previously thought to be colonized by only a few heterotrophic bacterial species. In addition, carbon and nitrogen cycling in these highly oligotrophic ecosystems are poorly understood. Here we genomically resolved a novel genus of *Thaumarchaeota, Ca. Nitrosodesertus*, found below boulders of the Atacama hyperarid core, and used comparative genomics to analyze their pangenome and site-specific adaptations. Their genomes contain genes for ammonia oxidation and the 3-hydroxypropionate/4-hydroxybutyrate carbon fixation pathway, indicating a chemolithoautotrophic lifestyle. *Ca. Nitrosodesertus* possesses the capacity for tolerating extensive environmental stress highlighted by the presence of genes against oxidative stress, DNA damage and genes for the formation of biofilms. These features are likely responsible for their dominance in samples with extremely low water content across three different boulder fields and eight different boulders. Genome-specific adaptations of the genomes included the presence of additional genes for UV resistance, heavy metal transporters, multiple types of ATP synthases, and divergent genes for aquaporins. Our results suggest that *Thaumarchaeota* mediate important carbon and nitrogen cycling in the hyperarid core of the Atacama and are part of its continuous and indigenous microbiome.

## Introduction

The surface soils in the hyperarid core (1) of the Atacama Desert are hostile environments characterized by extreme desiccation (water content < 1% by weight), high salt content resulting in low water activity, and high UV irradiation (∼ 30 J.m^-2^) (2). Scarce amounts of DNA from these soils have been analyzed in previous studies (2–4) revealing sparse microbial communities with low diversity, dominated by *Actinobacteria* and *Firmicutes*. While these previous studies showed that some of these microbes are likely alive and possibly active, as indicated by cultivation experiments (3) and *in-situ* replication measures [iRep; (5)] (2), very little is known about the carbon and nitrogen cycling in the hyperarid soils of the Atacama Desert. To date, only localized carbon fixation could be inferred from the findings of hypolithic and endolithic cyanobacteria (6,7), but no information on possible pathways for the transformation of other nutrients has been obtained so far. A recent study (8) of playas and alluvial fans located outside the hyperarid core reported the presence of Thaumarchaeal 16S rRNA sequences in the subsurface after a heavy rain event (40-90 mm) in 2015. However, without the genome-level information, Thaumarchaeal metabolic capability and contribution to the overall prokaryotic community could not be resolved in this study.

*Thaumarchaeota* mediate important environmental processes in both marine and terrestrial ecosystems and are particularly adapted to oligotrophic environments with their highly energy-efficient carbon fixation pathway (9,10). *Thaumarchaeota* have also been found in hot desert soils (e.g. Mojave Desert, California and Chihuahuan Desert, New Mexico) (11). However, most in-depth desert microbiome studies focus on bacterial communities (12,13) and multiple studies have reported decreasing archaeal diversity with increasing aridity (14,15). The general pattern of lower tolerance of *Archaea* to hyperaridity was supported by the absence of *Archaea* in other hyperarid environments such as the McMurdo Dry Valleys, Antarctica (12,16). The soil microbiome of the Atacama Desert has previously been thought to be dominated by *Bacteria*, with an exception of halophilic *Archaea* (*Halobacteriales*) in less arid locations such as coastal soils (2) and salt crusts (17). To our knowledge, previous molecular based studies of the hyperarid core revealed no evidence of *Archaea* and consequently, their adaptations and ecological roles in arid to hyperarid environments have not yet been studied.

The Atacama Desert hyperarid core harbors many expansive boulder fields (18–20) where individual boulders have been exposed for up to 37 millions of years (21,22) and transported by seismic activity. Despite the unique environments that the soils under the boulders present and the ubiquitousness of boulders in the Atacama Desert, no study has determined the microbial and geochemical composition below the boulders. In order to understand these uniquely protected hyperarid soils below the boulders that could harbor microbes playing a key role in carbon and nitrogen cycling in the Atacama Desert, we performed genome-resolved metagenomics of samples from the surface soil below the boulders and compared them to samples beside the boulders. Community structure and metabolic functions were interpreted in conjunction with geochemical measurements. *Thaumarchaeota* were revealed to be one of the key organisms differentiating microbial communities inhabiting below and beside boulders. Consequently, Thaumarchaeal genomes were selected for an in-depth pangenomic analysis to reveal adaptations to environmental stress and potential for carbon and nitrogen cycling. We investigated how *Thaumarchaeota* evolved in these uniquely protected, sparsely populated, and constantly selective environments.

## Material and methods

### Sampling location and procedure

Sampling was conducted in March 2019, in a dry period with the last recorded rain event occurring in June 2017 in the Yungay region. Three sampling sites, Yungay (Y), Maria Elena (M), and Lomas Bayas (L), were chosen based on a previous study (2) that identified inland hyperarid sites using the threshold of water content <1% by weight (**Figure 1a**). The coordinates of the three sample sites can be found in **Table S1**. Sampling was conducted in previously described characteristic boulder fields (18,19). At each boulder field, six boulders of diameter ∼50 cm and height ∼20 cm were chosen within a radius of ∼100 m from each other. For each boulder, two types of samples were taken, one below boulder (B) and one control sample (C) in the open soil ∼10 cm away from the boulder. All chosen boulders were well distanced from other boulders to make sure that the control samples were not constantly shadowed by other boulders or the sampled boulder itself. Samples were taken aseptically using precautions such as wearing a mask and sampling in upwind direction. New gloves were used for each sample, metal spatulas were previously autoclaved in aluminum foil, and newly unfoiled spatulas were used to scoop the topsoil (∼0.5 cm) into sterile 50 ml falcon tubes, which were then flash frozen in a liquid nitrogen dry shipper within half an hour of sampling. Control soil samples were taken first and then boulders were flipped over to sample below boulder soil as soon as possible to avoid aerial contamination. Additional samples were taken for geochemical analyses with a small shovel into a PE-sample bag (Whirl-Pak®, WI, USA) which were then stored at room temperature in the dark. See **Supplementary Materials and Methods M1** for additional field measurements.

**Figure 1.**
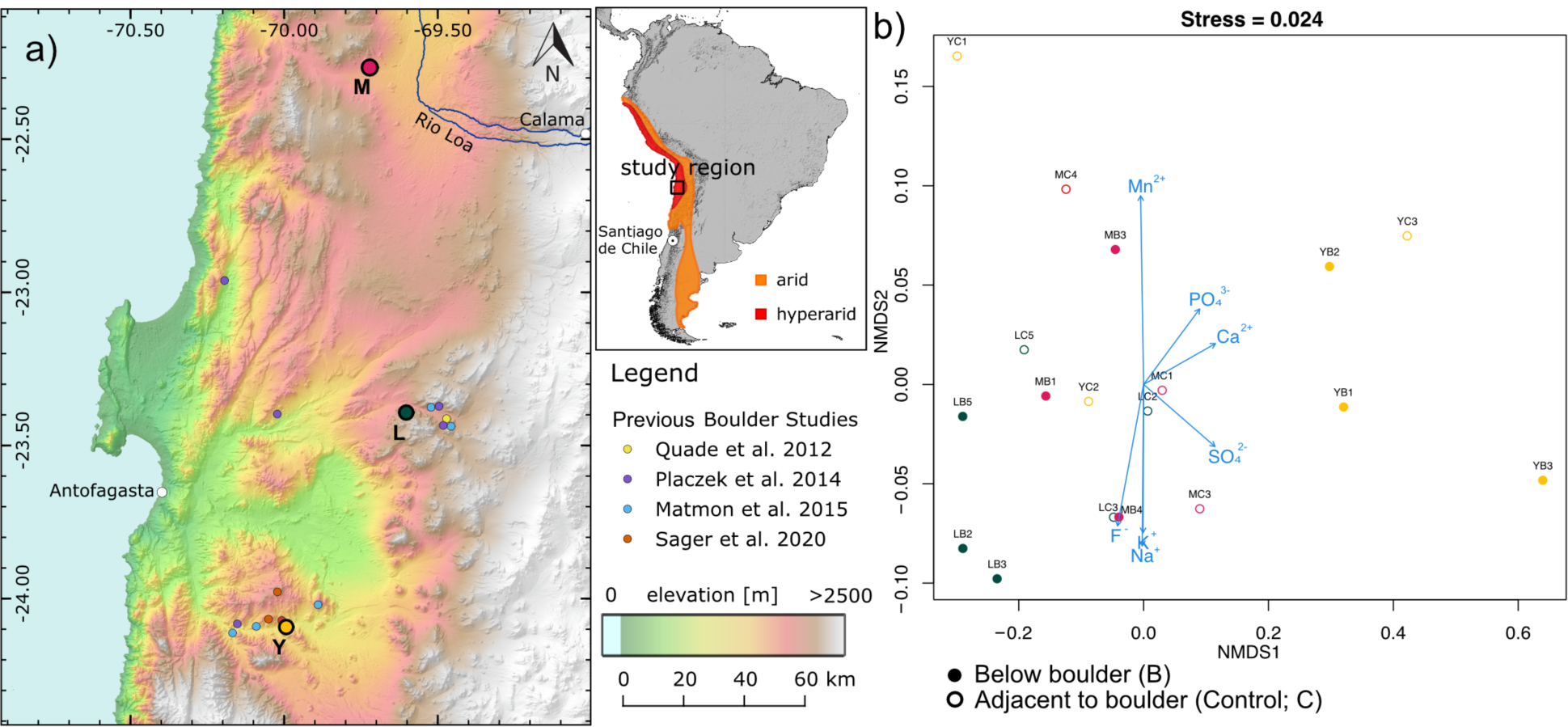
Sampling location and soil geochemistry. a) Location of three sampling sites and their abbreviations in parentheses b) Non-metric multidimensional scaling (NMDS) ordination of anion and cation concentrations in each soil sample. Different colors represent different sampling sites (Green = L, Red = M, Yellow = Y). Filled vs unfilled data points correspond to the sample type information. Blue vectors represent fitted ion species onto the ordination with a p-value less than 0.1.

### Geochemical and mineralogical analysis

Detailed methods for pH and electrical conductivity, anion and cation analysis, total organic carbon analysis and bulk mineralogy can be found in the Supplementary Materials and Methods M2-5.

***DNA extraction, Illumina library preparation and sequencing*** are presented in the **Supplementary Materials and Methods M6**.

### Metagenomic analysis, binning and annotation

P,Out of 24 attempted DNA extractions (**Table S2, Figure S1**), 15 yielded measurable amounts of DNA due to extremely low DNA content. Of those, eleven DNA extracts successfully yielded metagenomic libraries and subsequent metagenomic analyses were performed. For detailed methods on assembly, binning and annotation see **Supplementary Materials and Methods M7**.

#### Community analysis based on metagenomics

Operational taxonomic units (OTUs) were determined by extracting all S3 ribosomal proteins (rPS3) using hmmsearch (HMMER 3.2.1, http://hmmer.org/) across all assembled metagenomes. Retrieved rpS3 amino acid sequences were clustered using USEARCH (23) at 99% identity (24) and centroid sequences were extracted. Coverages of OTUs across all samples were calculated by mapping reads from each sample to the scaffolds of the centroids using Bowtie2 in sensitive mode (25) and filtering for a maximum of 5 mismatches (2% error rate) in both reads in each read pair. Coverages were then normalized by the total number of reads per sample. OTUs were placed into a phylogenetic tree by aligning using MUSCLE (26), alignment trimming using BMGE (BLOSUM30) (27) in default mode, and tree construction using iqtree v1.3.11.1 (28) with flags -m TEST -alrt 1000 -bb 1000. The phylogenetic tree was visualized using iToL (29). Shannon-Wiener Indices were calculated using the Vegan package (30) in R (31). ANOVA (34) analysis was conducted in R (31). Bray-Curtis (32) distance matrices were calculated for Principal coordinate analyses (PCoA), Non-metric multidimensional scaling (NMDS), BioENV(33), and Multiple response permutation procedures (MRPP, permutation=999) (35), which were subsequently visualized in R (31).

#### Phylogenomic analysis

Phylogenomic placements of the Thaumarchaeal metagenome-assembled genomes (MAGs) were determined using a supermatrix of 37 single-copy marker genes with all NCBI genomes annotated as *Thaumarchaeota* as of 4/6/2020 (36). The fact that the *Thaumarchaeota* classification on NCBI includes the newly reclassified phylum *Aigarchaeota* <u>(Hua et al. 2018)</u> allowed us to use the latter as an outgroup. CheckM (37) was used to quality filter genomes with thresholds <5% contamination, >50% completeness. Two local databases were created from the Atacama Desert and NCBI Thaumarchaeota MAGs (**Table S13**) respectively, against which homology searches were performed with HMMER 3.2.1 (http://hmmer.org/) using the HMM profiles for the Phylosift marker genes (36) with a cutoff of 1E-5. The resulting datasets were aligned with MUSCLE with default parameters (26) and curated manually to fuse contiguous fragmented sequences and remove extra gene copies. Ultimately, two genomes (GCA_011605725, GCA_011773305) were removed entirely, since they contained multiple sequences that were too distant from both *Thaumarchaeota* and *Aigarchaeota*. The resulting datasets were realigned as above, trimmed with BMGE (BLOSUM30) (27), and concatenated into a supermatrix of 312 operational taxonomic units (OTUs) and 7426 positions. Phylogenies were reconstructed with IQ-TREE 2 (28); first a tree with ModelFinder (38) (-m MFP -bb 1000 -alrt 1000 -abayes) that served as guide tree for a run with the PMSF model (39) (-m LG+C60+F+G -bb 1000 -alrt 1000 -abayes). As per the suggestion of the IQ-TREE authors, we considered those branches strongly supported with at least 95 for ultrafast bootstrap (40) and 80 for the SH-aLRT test (41).

#### Comparative genomics

The predicted protein sequences of eight NCBI *Ca. Nitrosocosmicus* reference genomes (**Table S3**) were compared with the recovered Thaumarchaeal MAGs. The CompareM package (github.com/dparks1134/CompareM) was used to identify the orthologous fraction (OF) and calculate the average amino acid identity (AAI) of orthologous genes between a pair of genomes, and fastANI (42) was used to calculate the average nucleotide identity between genomes using default parameters. OrthoVenn2 (43) was used to identify and visualize orthologous clusters across genomes.

## Data availability

All sequencing data will be submitted to SRA and genomes will be made publicly available through NCBI.

## Results and Discussion

### Hyperarid soils sheltered under the boulders are geochemically distinct and organic carbon deficient

As previously documented (18–20,44), a substantial part of the Atacama Desert hyperarid core features expansive boulder fields, where the topsoil is covered by boulder-sized clasts (**Figure. 1, Figure S4**). Soils below the boulders experience lower diurnal temperature and relative humidity fluctuations than soils beside the boulders (**Figure S2a-c**). Based on the dew point temperature calculations, we showed that the condensation of water in the morning hours is far less likely for soil below boulders compared to soil beside boulders (**Figure S2d-f**), suggesting that water content below boulders may be even lower than previously studied Atacama Desert hyperarid top soils (∼0.2% by weight) (2).

We compared soil samples of two sample types: soils taken below boulders (B) and soils taken adjacent to the boulder (control, C), at three different sampling locations (Lomas Bayas, L; Maria Elena, M; Yungay Valley, Y) (**Figure 1a**). While the collected soils were mineralogically very similar with some variation with sampling location (**Figure S3**), their ion concentrations showed large variance between boulder fields, individual boulders, and sample types. Interestingly, B samples clustered based on their sampling location, while C clustered independent of sampling sites (**Figure 1b)**. In general, samples from locations L and M were enriched in F^-^, while Y samples were enriched in PO_4_^3-^, SO_4_^2-^, Mn^2+^ and Ca^2+^, suggesting boulder field specific patterns of ion concentrations. More sampling location dependent ion composition patterns amongst the B samples indicate that soils below the boulder are sheltered from external input of ion species (i.e. atmospheric deposition), thereby exhibiting a more representative ion composition patterns of the soils in the area beyond the topsoil. When comparing the B and C sample of each individual boulder, nitrate and magnesium ion concentrations were significantly lower in B samples compared to C samples (paired t-test; NO_3_^-^: t(8) = -3.9, p-value = 0.0451, Mg^2+^: t(8) = -2.33, p-value = 0.0484, **Figure S5**). Total Organic Carbon (TOC) concentrations were at or below detectable levels (**Figure S6**) in both below boulder and beside boulder samples. Our results show that the soils below the boulders are not only hyperarid and organic carbon deficient, but also sheltered from the atmospheric input of both water (e.g. fog, dew) and ion species.

### An actively replicating microbial community in the Atacama Desert hyperarid core

We investigated the eleven successfully prepared metagenomes (for details see Material and Methods and **Table S4**): three below boulder and three control samples came from Lomas Bayas (LB2, LB3, LB5 and LC2, LC3, LC5), three samples below boulder were from Maria Elena (MB1, MB3, MB4) and two below boulder samples from Yungay Valley (YB1, YB3).

Genome-resolved metagenomics of these eleven samples yielded 73 high quality (>75% completeness, <15% contamination) metagenome-assembled genomes (MAGs), of which 71 belonged to only three different phyla, reflecting the limited diversity of this extreme ecosystem. Eight of these high quality genomes were Thaumarchaeal with completeness ranging from 84.67 to 98.54% and contamination below 10% (**Table S4**). Other high quality MAGs belonged to *Actinobacteria* (n=34), *Chloroflexi* (n=29), *Firmicutes* (n=1), *Alphaproteobacteria* (n=1). *In situ* replication measures [iRep, (5)] were successfully calculated for 32 out of all high-quality bacterial genomes (n=65), indicating an active metabolism of the majority of the indexed population (calculated iRep values of 32 genomes ranged between 1.34 and 3.47, mean: 1.98). On average, genomes recovered from below boulder metagenomes were associated with higher iRep values than the control metagenomes (p-value < 0.04, Welch’s t-test, **Figure S7)**. A full overview of genome statistics, their taxonomic classification and corresponding iRep values is provided in **Table S4**.

### Atacama soils below boulders harbor unique microbial communities with high shares of Thaumarchaeota

We detected 147 different *Bacteria* and *Archaea* based on clustering of S3 ribosomal proteins (rpS3, 99% identity, **Table S5, Figure S8**). Shannon indices showed higher variation in alpha diversity for below boulder samples compared to control samples (**Figure S9**). Principal Coordinate Analysis (PCoA) of the communities (**Figure S10a**) demonstrated clustering of samples based on the sample site (L, M, Y) as well as the sample type (B, C). This was corroborated by the Multiple Response Permutation Procedure (MRPP) indicating significant influence on the community structure by both sampling location (chance corrected within group agreement A = 0.2648, significance of delta = 0.001) and sample type (A = 0.1488, significance of delta = 0.002). Using BioENV (33), we identified F^-^concentration to be most correlated (correlation = 0.573) with the community composition. Additionally, we conducted a NMDS analysis (**Figure S10b**), identifying additional ions (Ca^2+^, SO_4_^2-^, K^+^, Cl^-^) that could be correlated with the community composition.

Out of the 147 different taxa, 33 were identified to be significantly different in their abundances (ANOVA (34) p-value < 0.05) between B and C samples (**Table S6**). Such taxa included *Actinobacteria* (belonging to *Cryptosporangiaceae, Streptomycetaceae*, and *Geodermatophilacaea*), as well as one *Alphaproteobacteria* (*Acetobacteraceae*). These taxa were particularly abundant in control samples and near absent in below boulder samples, suggesting specific and unknown selection processes for the two different sample types. Alternatively, some of these taxa may be deposited through aeolian transport (45). **Figure 2** shows the phylogenetic relationship between the top 30 most abundant taxa across the samples based on rpS3 proteins and links them to their respective MAGs as well as their differential coverage across the samples. We conclude that below boulder (B) and beside boulder (C) present substantially different habitats of the same ecosystem.

**Figure 2.**
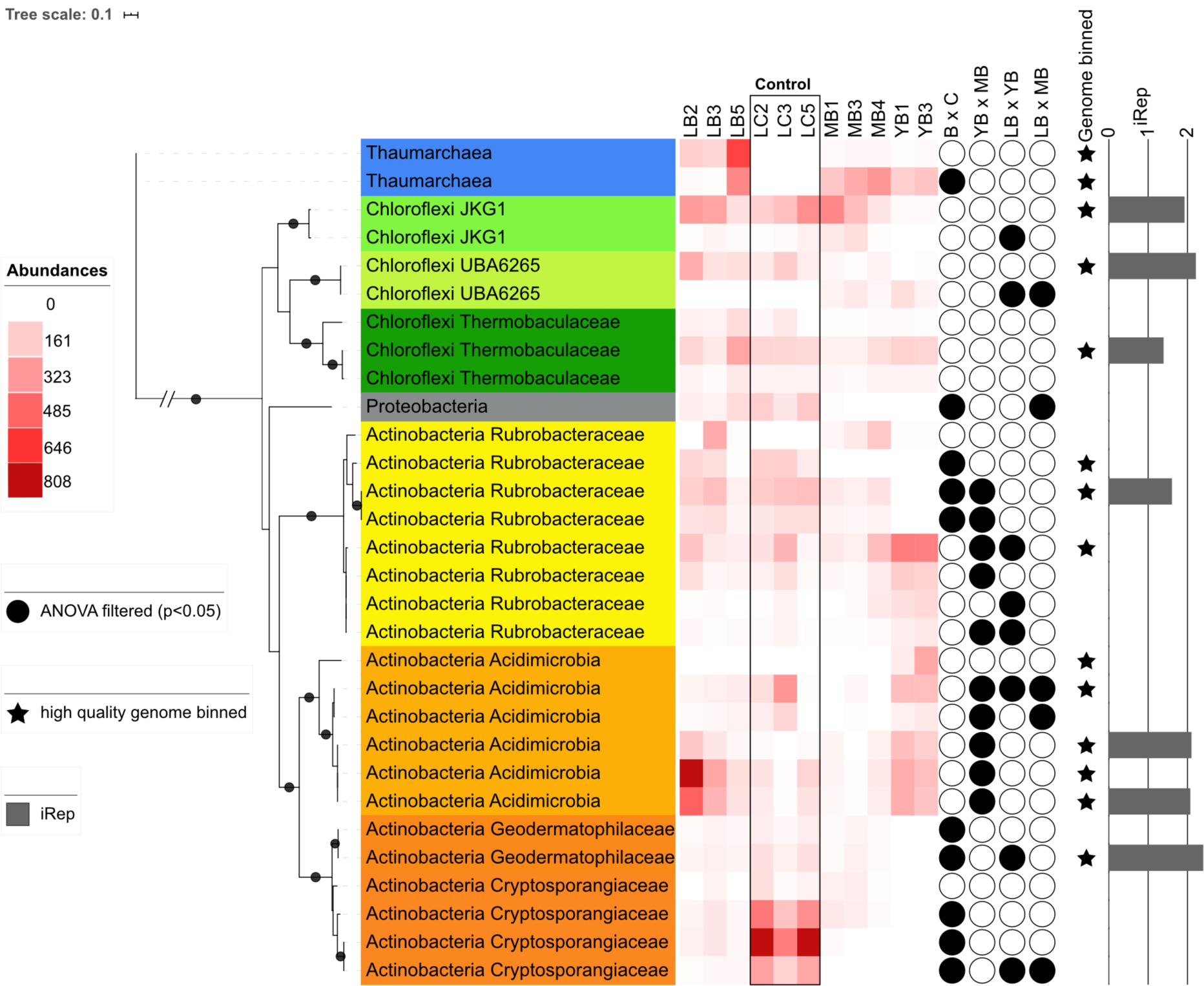
Phylogenetic tree of 30 most abundant taxa (rps3 clusters) out of 147 and their normalized abundances across all samples. Filled stars represent successful binning of the OTU in high quality genomes and red bars indicate iRep values calculated for the high quality genomes. Strongly supported branches as described in the M&M section are indicated with black dots.

One *Thaumarchaeal* OTU was the only taxon based on ANOVA (34) (p-value = 0.0396) with a higher abundance below boulders and near absence in control samples. All eight below boulder metagenomes contained high abundances of *Thaumarchaeota*. Based on the ranked abundance of ribosomal protein S3 (rpS3) gene coverages, *Thaumarchaeota* ranked amongst the top seven most abundant taxa across all below boulder samples. In three samples (MB3, MB4, LB5), *Thaumarchaeota* were the most abundant organisms, e.g. in LB5, *Thaumarchaeota* were 4-fold more abundant than the second most abundant taxon. The abundance of *Thaumarchaeota* under boulders and their near absence in the nearby irradiated soil support the previous findings from marine environments (10,46) where surface waters harbored lower abundances of *Thaumarchaeota*. Photoinhibition of ammonia oxidation in ammonia oxidizing archaea (AOA) (47) has previously been hypothesized as the cause, along with other proposed hypotheses, such as increased competition (48,49) and indirect photoinhibition by Reactive Oxygen Species (ROS), such as hydrogen peroxide (50). To date, the underlying reason for the lower abundance of *Thaumarchaeota* in highly irradiated environments remains inconclusive. Based on the low amounts of DNA recovered from most control samples (< 1.14 ng / g soil), we conclude that the near absence of *Thaumarchaeota* in the open irradiated top soils is likely not due to increased competition, at least at our study sites. Photochemically produced ROS (H_2_O_2_ and metal superoxides and peroxides) have previously been found to accumulate in the Atacama Desert (Yungay site) top soils at levels an order of magnitude higher than in non-arid control soils (51). Additionally, in contrast to ocean environments, UV and photoradiation do not penetrate into the soil beyond the very surface of the topsoil and with minimal soil turbation in the Atacama, we expect the effect of UV and photoradiation inhibition reaching below the top ∼ 0.5 cm of soil unlikely. Therefore, we hypothesize the inhibition of high ROS levels (50) to be the main reason why *Thaumarchaeota* are not abundant in the control samples despite their potential desiccation tolerance.

### Ammonia Oxidizing Thaumarchaeota occupy an important niche in the Atacama Desert carbon and nitrogen cycling

Normalized abundance of key marker genes in the assembled metagenomes revealed potential for C1 metabolism (**Figure 3, Table S7-8**), complex carbon degradation, and fermentation across all samples. Three carbon fixation pathways (3-hydroxypropionate cycle, 3HP/4HB, CBB cycle) were detected however, abundances of each marker gene varied with site. No 3HP/4HB cycle was found in control samples, while the 3-hydroxypropionate cycle was found in low abundance in MB and YB samples. Significant gaps in the potential for nitrogen cycling were observed. Nitrile hydratases were found across all samples, and archaeal ammonia oxidation potential was only found in below boulder samples (with the exception of YB1). No potential for nitrogen fixation, nitrate-, and nitric oxide reduction as well as nitrite oxidation were identified. The lack of nitrogen fixation and denitrification genes suggest low overall biological input and little loss of biologically available nitrogen. Although the investigated soils are known to be enriched in nitrates (particularly at ∼1m depth) that have accumulated over millions of years through abiotic processes (e.g. atmospheric formation through lightning followed by dry deposition and rainwater infiltration) (52), below boulder nitrate concentrations are significantly lower (**Figure S5**), likely due a combined effects of highly microbial activity and lack of atmospheric or hydrologic input. Therefore, we propose that nitrogen cycling below boulders is largely controlled by microbial activity. Specifically, we suggest a highly equilibrated nitrogen cycle with *Thaumarchaeota* nitrifying ammonia produced through protein ammonification performed by diverse *Chloroflexi* and *Actinobacteria* (**Table S9**) in these below boulder environments.

**Figure 3.**
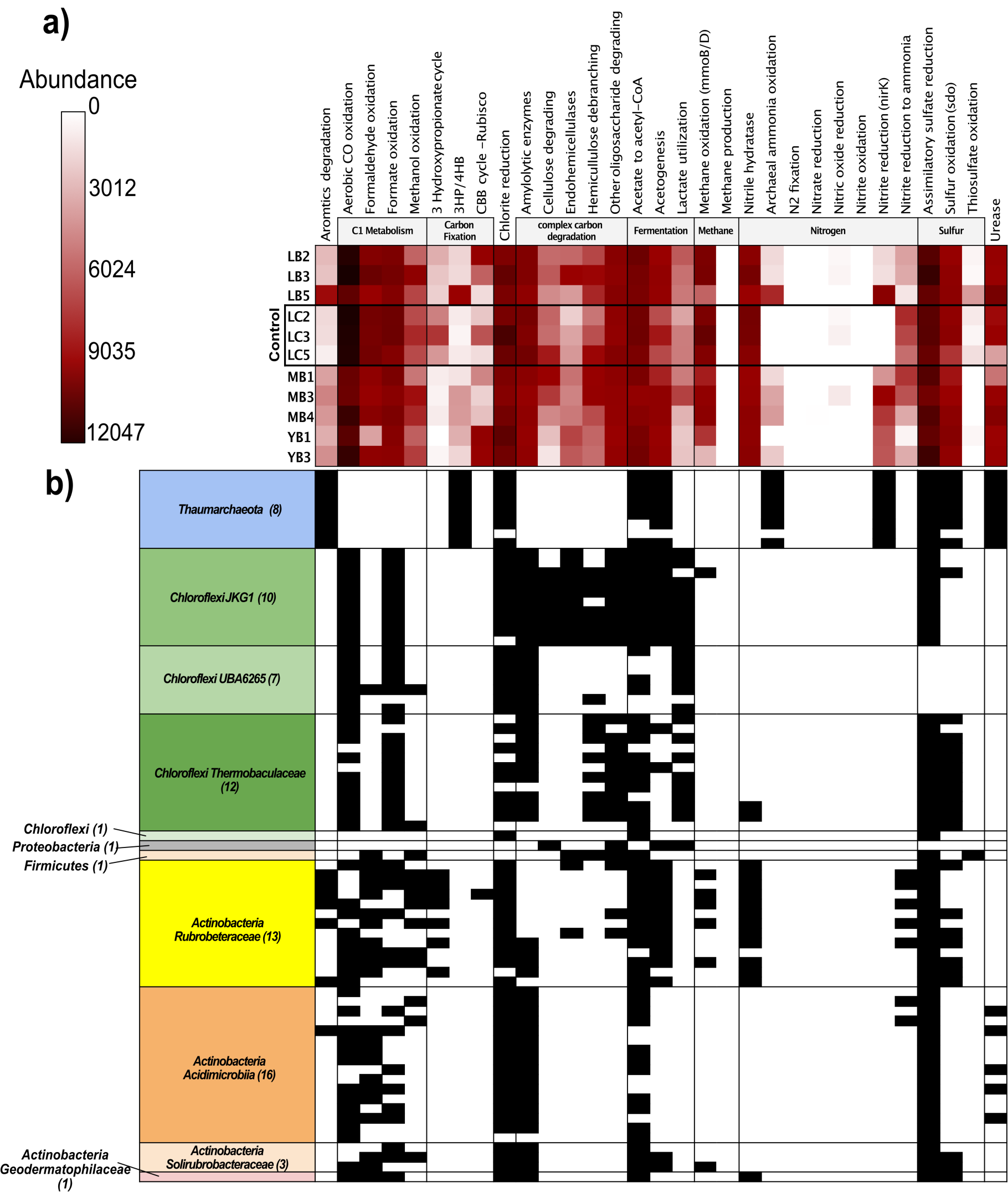
Metabolic potential prediction across samples and high quality genomes. a) Normalized abundances of chemoautolithotrophic marker genes predicted using METABOLIC for each sample. b) Presence (black) and absence (white) of chemoautolithotropic marker genes in high quality genomes. Genomes are clustered based on taxa and the number of genomes in each cluster is shown in parentheses in the row names.

Resolving the metabolic potential at the genomic level delineated the role each taxonomic group plays in this highly streamlined community. Presence and absence of key metabolic genes for each high quality genome are shown in **Figure 3b**. Our analysis shows that although all samples show carbon fixation potential, taxa capable of fixing carbon are limited to *Thaumarchaeota* (through the 3HP/4HB pathway) and some *Rubrobacteraceae* (3-hydroxypropionate pathway). Surprisingly, only one Form I Rubisco could be binned to a high quality *Rubrobacteraceae* genome, despite a stronger signal seen in the metagenomes. Chloroflexi genomes associated with the lineage *JKG1* had the broadest potential of degrading complex carbon and were capable of a fermentative lifestyle, while *Actinobacteria* could metabolize a wider range of C1 substrates. Nitrite reduction potential detected in below boulder sites was constrained to the nirK genes found in *Thaumarchaeota*. NirK in *Thaumarchaeota* has been hypothesized to play a key role in ammonia oxidation (53), and is biochemically capable of transforming N compounds to produce nitric oxide (54). However, whether it also denitrifies organic nitrite leading to a loss of organic N in a natural environment remains to be confirmed. Genome-resolved metabolic predictions revealed conserved metabolic capacities across genomes that belong in the same taxonomic family, with *Thaumarchaeota* playing a unique role in the nitrogen and carbon cycling in the Atacama hyperarid core.

### A novel genus of Thaumarchaeota with highly conserved core genome and diverse auxiliary genes

Eight high quality Atacama Boulder Thaumarchaeal genomes (ABT) were assembled with an average GC content of 34.6% (± 0.1%) and average size of 2.5 Mbps (± 0.4 Mbps). Each genome contained on average 3,123 (± 579.7) predicted genes with a mean coding density of 71.8% (± 1.7%). The genomes were phylogenetically placed using 37 single-copy house-keeping genes, forming a monophyletic sister cluster to the recently characterized *Ca. Nitrosocosmicus* (**Figure 4a**). The ABT clade and *Ca. Nitrosocosmici* form a sister group to *Ca. Nitrososphaera*, a mesophilic terrestrial clade. Genomes from the same sites were more related to each other, with ABT-MB and ABT-YB genomes forming a separate branch from the ABT-LB genomes (**Figure 4b**). One copy of the ammonia monooxygenase A (*amoA*) gene was found in five ABT genomes (**Table S10**). Upon closer look, two other genomes (ABT-MB3, ABT-MB4) contained conserved *amoA* regions with unresolved assembly errors and therefore failed in protein prediction. No *amoA* sequences were found in ABT-YB1. 3 additional unbinned *amoA* genes were detected across the metagenomes (MB3, MB4, YB3). Altogether, the eight *amoA* nucleotide sequences were 100% identical in their amino acid sequences to each other and to previously published *amoA* sequences from *Ca. Nitrosocosmicus oleophilus* and *Ca. Nitrosocosmicus exaquare*, which had been phylogenetically identified to be one of the basal clades of archaeal *amoA* after *Ca. Nitrosocaldus (55)*. **Figure S11** resolves the nucleotide level phylogenetic placement of binned *amoA* sequences as well as unbinned *amoA* sequences recovered from the sample metagenomes. Interestingly, one *amoA* recovered from a low quality bin (68% completeness; 5.8% contamination) in the YB3 metagenome (node “ABT-YB3 (low quality bin)” in **fig. S11**) was divergent (∼80% ID) from the rest at the nucleotide level, while 95.8% identical to other ABT and *Ca. Nitrosocosmicus amoA* genes at the amino acid level. The rpS3 gene recovered from this bin was classified as Thaumarchaeal, with 75% identity to other binned rpS3 in ABT, and its closest NCBI reference sequence being *Ca. Nitrosocosmicus* sequences at 65% identity. This divergent Thaumarchaeal bin was approximately three-fold less abundant than another Thaumarchaeal bin (ABT-YB3) recovered at a higher quality from the same metagenome (YB3). Due to low quality and lower abundance of this divergent bin, our study focuses on other eight high quality genomes that are much more closely related and found across all metagenomes under the boulder including YB3.

**Figure 4.**
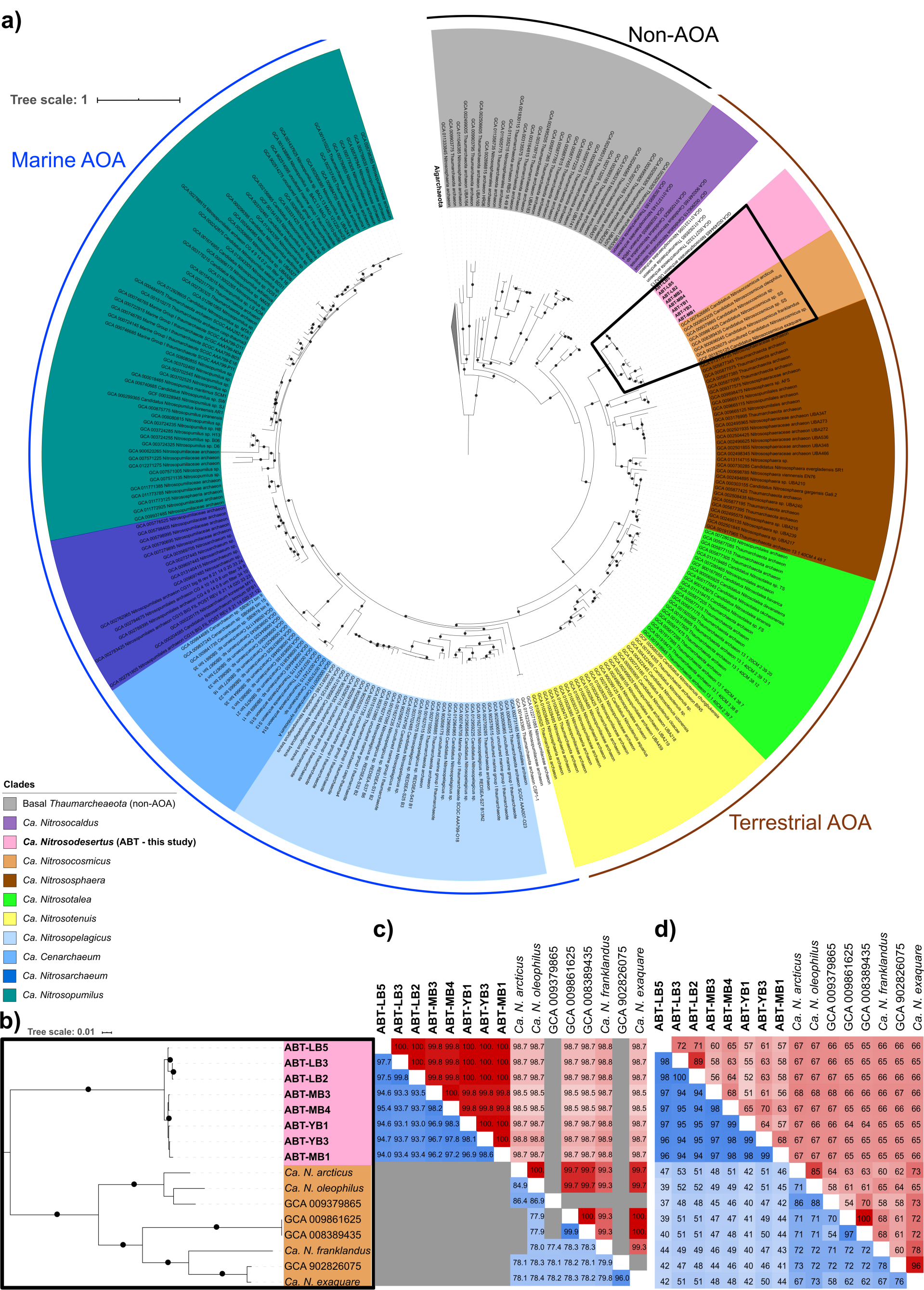
Phylogenomic placement of ABT genomes using 37 housekeeping single-copy genes. a) Phylogenetic tree of 298 NCBI genomes annotated as *Thaumarchaeota* and 8 ABT genomes. *Aigarchaeota* were identified and used as the outgroup. Black, brown and blue ranges distinguish whether organisms are Ammonia Oxidizing Archaea (AOA) and their typical habitats (terrestrial vs marine). Strongly supported branches as described in the M&M section are indicated with black dots. b) Zoomed view of the branches placing the ABT genomes and its sister group *Ca. Nitrosocosmicus*. Strongly supported branches as described in the M&M section are indicated with black dots. c) Lower-right (blue) triangle of the matrix corresponds to FastANI between genomes, where gray values indicate below calculation threshold (80% identity). Upper right (red) triangle of the matrix corresponds to 16S rRNA identity values, where gray values are used for genomic bins without a 16S rRNA. d) Lower right (blue) triangle corresponds to the Amino Acid Identity (AAI) and upper right (red) triangle corresponds to the Orthologous fraction (OF) between a pair of compared genomes.

In order to taxonomically resolve the eight recovered *Thaumarchaeota* genomes, we compared them to *Ca. Nitrosocosmicus* genomes that had been isolated or metagenomically assembled from around the world (**Table S4**) in diverse environments ranging from the arctic soil (56), tar-contaminated soil (57), vegetable field (58), dinosaur fossil (59) to wastewater filters (60). High ANI (93.0 - 99.8%) (**Figure 4c**) between ABT genomes indicated that all ABT genomes belong to one genus. Using the ANI threshold of 95% (61,62) for species delineation, we identified two species within the ABT clade, with genomes recovered from LB site belonging to one species and the rest to another. The mean Amino Acid Identity (AAI) of 53.9% between pairs of *Ca. Nitrosocosmicus* and ABT genomes (**Figure 4d**) fall below the genus delineation threshold of 65% (63) indicating that the two clades form separate genera. Based on these findings, we propose two new species names that belong to a new genus: *Ca. Nitrosodesertus atacamaensis* (ABT-LB2, ABT-LB3, ABT-LB) and *Ca. Nitrosodesertus subpetramus* (ABT-MB1, ABT-MB3, ABT-MB4, ABT-YB1, ABT-YB3).

While the eight ABT genomes share a highly conserved core genome (mean AAI = 96.5 %), between 11% and 49% (mean = 37.7%) of the genes had no other orthologs in the recovered genomes despite the relatively similar and static environmental conditions that they were found in. High AAI in the orthologous fraction of the eight ABT genomes and conserved *amoA*s recovered in sites more than 200 km apart from each other suggest that ABTs originated from the same strain of *Thaumarchaeota*. However, large fractions of sample-specific auxiliary and divergent genes suggest site-specific adaptations to their respective isolated habitats.

### Pangenomic comparison of ABT genomes and their sister clade reveal unique adaptations including heavy metal resistance, biofilm formation, water transport and sodium bioenergetics

In order to understand the conserved metabolic potentials between ABT and *Ca. Nitrosocosmicus*, unique adaptations of the ABT in the Atacama Desert, and niche differentiations between sites, we analyzed the highest quality (> 95% completeness, <5% contamination) genomes (ABT-LB3, ABT-MB4, ABT-YB3) from each of the three sites along with three (near)-complete *Ca. Nitrosocosmicus* reference genomes (*Ca. N. franklandus, Ca. N. oleophilus, Ca. N*.*exaquare*). 1287 homolog clusters are shared across all six genomes (**Figure 5, Table S10**). For example, all genomes contained a highly conserved AmoABX operon, although only two out of eight ABT bins contained 1-2 *amoC* copies. All genomes revealed the metabolic potential for mixotrophy along with important genes for nitrogen cycling, including genes for copper-dependent nitrite reductase (*nirK*), urease (*Ure*), urea transporter, ammonium transporter, deaminases, lyases, and carbonic anhydrase (**Table S11**). Additionally, amongst the shared genes we found key stress response genes that could provide resilience against highly oxidizing environments (**Table S10**).

**Figure 5.**
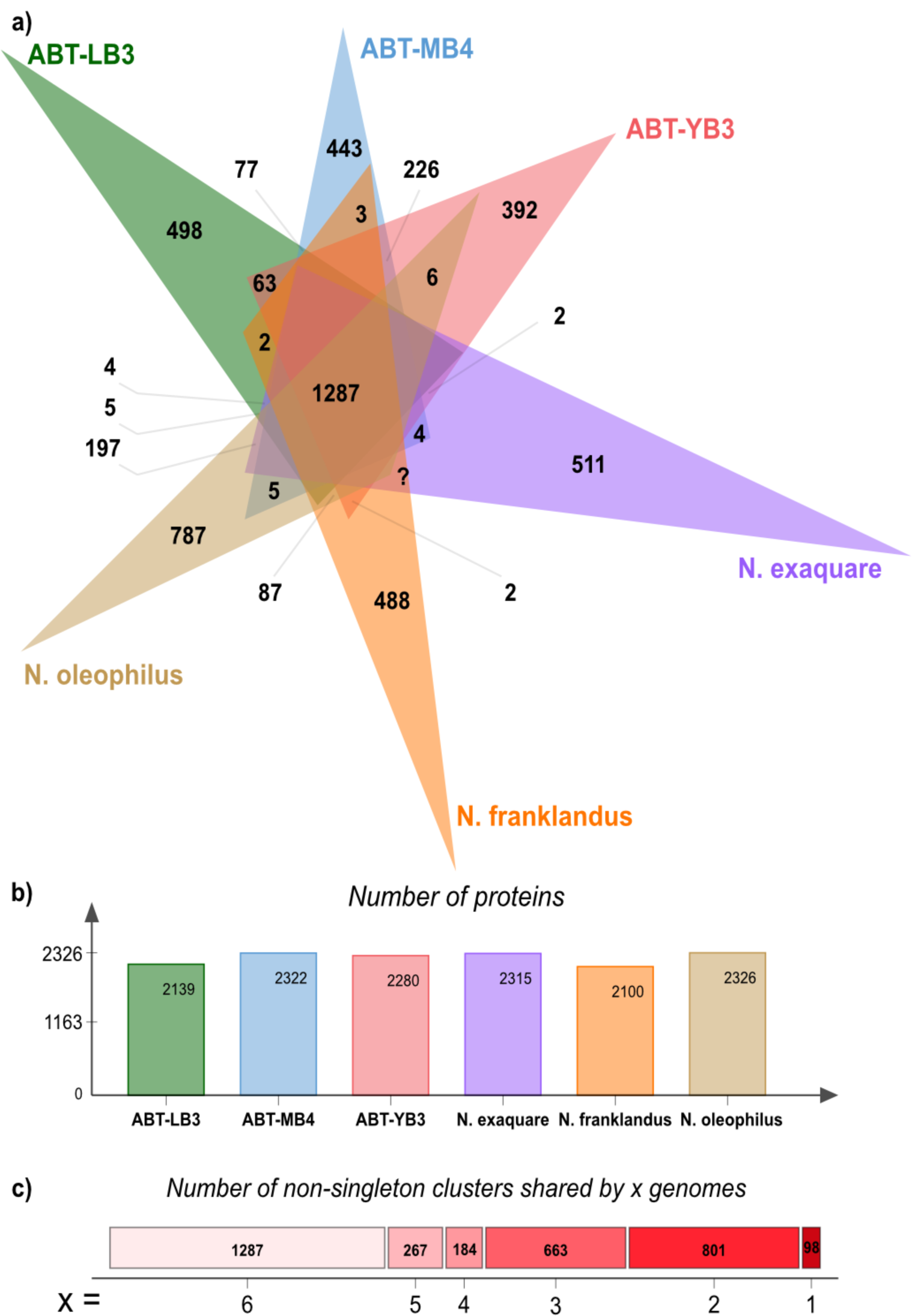
Shared and auxiliary protein clusters of ABT and its sister genus. a) Shared orthologous protein clusters (including singletons) across six genomes (three Ca. *Nitrosocosmicus*, three Ca. *Nitrosodesertus* [ABT]). b) Number of proteins in each genome. c) Number of orthologous protein clusters (excluding singletons) shared across x number of genomes.

296 protein clusters were shared between ABT genomes but were not present in *Ca. Nitrosocosmicus* genomes (**Figure 5)**. Notable genes identified in these clusters include those involved in biofilm production and cell adhesion capacity (**Table S10**). The ability of Ca. *Nitrosocosmicus oleophilus* to form biofilms and produce exopolysaccharide (EPS) has previously been demonstrated by Jung *et al*. (57). EPS production and biofilm formation in general are considered major adaptation mechanisms for xerotolerant bacteria (64) and ABT genomes may also employ this mechanism to protect against desiccation.

12.8% to 16.5% of the protein coding genes in each ABT genome belonged to unique protein clusters or were singletons (**Figure 5, Table S12)** that did not share any similarity to other genes in the ABT genomes. Amongst singletons of the genome ABT-LB3, many were involved in membrane transport of metals, such as magnesium, copper and cobalt transporters as well as lead, cadmium, zinc and mercury transporting ATPases, potassium uptake proteins and bacterioferritin. The presence of these genes may be an adaptation to heavy metals known to accumulate in the Atacama Desert soils (65). Similarly, notable singletons found in genomes ABT-MB4 and ABT-YB3 included putative cobalt transporter, fluoride transporters, zinc uptake system, mercuric reductase, ferrous iron permease, and phosphite transport system. See supplementary results for further findings from the pangenome analysis.

Further genome comparisons across all eight ABT genomes revealed additional key adaptations to desiccation and osmotic stress. We identified up to seven different copies of water channel membrane proteins (aquaporin Z2) (66) per genome. Interestingly, some of these proteins were highly divergent from each other at the AA sequence level, while others were truncated (**Figure S12**) despite being found mid-contig with relatively conserved surrounding genes. Multiple copies of aquaporin genes per genome as well as the divergent and truncated subset indicate possible genome specific adaptations to desiccation and osmotic stress. We also recovered two distinct types of ATP synthases (namely A-type and V-type (67,68)) from the eight ABT genomes. Three ABT genomes (ABT-LB2, ABT-LB3, ABT-YB1) contained only A-type ATP synthase, while the rest contained both the A-type and the V-type ATP synthases often in multiple copies. Wang *et al*. (67) concluded that the V-type ATP synthases are horizontally transferred from *Euryarchaeota* and conserved among the acidophilic and hadopelagic *Thaumarchaeota*, potentially playing a key role in adaptations to acidic environments and elevated pressure through proton extrusion. Considering that Atacama Desert soils are slightly alkaline (average pH = 7.7 **Figure S13**), it is surprising that the V-type ATP synthase is found and conserved across five ABT genomes. Zhong *et al*. (68) hypothesized that these V-type ATP synthases may be coupled with Na^+^motive force instead of proton pumping. Atacama Desert soils present high salt stress, and therefore the V-type ATP synthase could perform Na^+^pumping and provide protection against high sodium stress (**Table S14**). Notably, all genomes featured high-affinity Na^+^/H^+^antiporter NhaS, with ABT-LB2 and ABT-LB3 genomes featuring five copies, while the others featured a single copy. This may be correlated to the lack of Na^+^binding V-type ATP synthase in ABT-LB2 and ABT-LB3 genomes. Additional genes associated with Na^+^bioenergetics were identified, including sodium/glucose transporter, putative calcium/sodium:proton antiporter, sodium bile acid symporter family protein, sodium/hydrogen exchanger and sodium-dependent dicarboxylate transporters. This suggests that ABT genomes are not only highly adapted to high salt concentrations but also are potentially capable of utilizing the sodium gradient to scavenge useful biomolecules for mixotrophic growth as well as generate ATP in the hyperarid core of the Atacama Desert.

## Conclusions

We report here the first evidence of highly adapted ammonia-oxidizing *Thaumarchaeota* inhabiting the hyperarid Atacama Desert in high relative abundance, including the first systematic comparison of microbial communities found below boulders of the Atacama Desert hyperarid core with the microbial communities present in the open, unprotected desert soil. This study expands the realm of Thaumarchaeal presence revealing high adaptability and resilience to hyperarid, high salt and low-nutrient environments. In-depth genomic characterization of these ABT genomes elucidated their niche potential roles in N and C cycling in highly nutrient deficient Atacama Desert soils, as well as key adaptations against oxidative stress, salt stress and hyperaridity. By comparing the eight closely related ABT genomes retrieved from these isolated and disconnected habitats, we hypothesize *Ca. Nitrosodesertus* to be a potentially endemic *Thaumarchaeota* genus in the Atacama Desert, with organisms in this genus harboring highly conserved shared genes and large numbers of site-specific auxiliary genes. Beyond the Atacama Desert, this study provides a blueprint for future studies of extreme terrestrial environments (i.e. Antarctic and extraterrestrial) where finding pockets of pristine, sheltered and contained environments, as simple as below boulders, could lead to a discovery of uniquely conserved communities and help delineate the indigenous microbial community members adapted to extreme conditions.

## Acknowledgements

This work was funded by ERC Advanced Grant HOME (# 339231) to DSM. AJP and TVLB were supported by the Ministerium für Kultur und Wissenschaft des Landes Nordrhein-Westfalen (“Nachwuchsgruppe Dr. Alexander Probst”). We thank Bärbel Försel for insightful discussions, Manuela Alt and Kirstin Weiß for TOC measurements, Thomas Neumann for providing access to the XRD laboratory, and Iris Pieper and Claudia Kuntz for technical assistance and measurement of water-soluble ion species.

## Competing Interests

All authors declare that they have no competing interests.

## Author contributions

DSM and YH conceived the project; YH, FLA, and AA planned and conducted sampling; JSS and MS prepared metagenomic libraries and performed sequencing as well as initial quality filtering of reads; YH assembled, curated and analyzed sequence data with contribution from TLVB and AJP; AJP provided computational resources; FLA performed geochemical analyses and AA provided input in data interpretation; PSA performed phylogenetic analysis; YH wrote the manuscript with contribution from AJP; all authors discussed and revised the manuscript.

## Supplementary Information

### List of Content

Supplementary Materials and Methods (M1-M7)

Supplementary Results and Discussion

Figures S1-S12 and Tables S1 and S2

Legends for Supplementary Tables S3-14

Descriptions for additional Supplementary File

## Supplementary Materials and Methods

### M1. Field measurements

The HOBO U23 pro Temperature/Relative Humidity data logger (Onset, Cat# U23-001, MA, USA) was used to monitor the temperature and relative humidity of each site at the time of sampling. For each sampling site (Y, M, L), an extra boulder was chosen for conducting HOBO logger measurements. One logger was placed under a boulder similarly sized to those chosen for sampling, and another logger was placed ∼20 cm away from the boulder on the open soil. For the Y site, a continuous measurement over 130 days (15 March - 25 July 2019) was conducted for characterizing diurnal fluctuations for both below boulder, beside boulder and 1 m above ground. Logged data was then used to calculate dew point temperatures as described in Lawrence *et al. (1)*.

### M2. pH and electrical conductivity

To evaluate the pH and the electric conductivity (EC) of the soil, samples were prepared in a ratio 1:5 v/v (5 ml sample to 25 ml distilled water), shaken for one hour to prevent the particle from settling and sedimented for another hour, before measuring pH (691 pH Meter, Metrohm, Switzerland). The standard deviation was determined by repeated measurements of in-house standards, SD = 0.24 (n = 16). EC was measured with a handheld electric conductivity meter (GMH 3400, Greisinger, Germany). Reproducibility variation was 5% (n = 3). Both measurements were conducted at the Center of Astronomy and Astrophysics the Technische Universität Berlin.

### M3. Anion and cation analysis

Samples and processing controls for water-soluble ion analysis were prepared based on the standard DIN EN 12457 - 4 (2003) protocol. Briefly, samples were sieved to obtain <2 mm particles which were used to prepare an eluate of a 1:10 w/w (4.5 g sample to 45 g distilled water). After 24 h of continuous shaking, the eluate was filtered through 0.2 µm mesh and stored at -20°C until measurement. Anionic species (Cl^-^, NO_3_^-^, PO_4_^3-^, SO_4_^2-^) were measured by ion chromatography (DIONEX DX-120 Ion chromatograph, Thermo Fisher Scientific, USA, with a guard column AG 22, 4×50 mm and an analytical column AS 22, 4×250 mm). Reproducibility variation was <1% (n = 5). Cations (Ca^2+^, Fe^2+^, K^+^, Mg^2+^, Mn^2+^, Na^+^) were determined by inductively-coupled plasma optical emission spectrometry (iCAP 6000 ICP Spectrometer, Thermo Fisher Scientific, USA). Reproducibility variation was <5% (n = 5). Both analyses were conducted at the Department of Soil Science of the Technische Universität Berlin. Bray-Curtis distance (“vegan” package (2)) metric was used to calculate the distance matrix of samples based on the ion concentrations, which was then used for non-metric multidimensional scaling analysis in R (3).

### M4. Total organic carbon analysis

The total organic carbon (TOC) was measured with an elemental analyzer (Vario Max C, Elementar, Germany) using catalytic tube combustion at the Department of Life Science of the Humboldt Universität Berlin. Samples were first ground to powder. Due to low TOC concentrations, 1 g was used for combustion. At 600°C the organic carbon was removed under the carrier gas nitrogen and oxidized by oxygen in the presence of copper oxide. Remaining elemental carbon was combusted with the addition of oxygen. The resulting CO_2_ was then determined successively by infrared detection. The measurement was conducted in duplicates with a detection limit of 0.0124 wt%.

### M5. Bulk mineralogy

For the bulk mineralogy, soil samples were homogenized and ground to powder. X-ray powder diffraction (XRD) analysis of the soil salts was performed by using a powder diffractometer (D2 Phaser, Bruker, USA) at the Department of Applied Geochemistry of the Technische Universität Berlin. The X-ray source was Cu Kα radiation (K-alpha1 = 1.540598 Å, K-alpha2 =1.54439 Å) with a performance of 30 kV and 10 mA. A step interval of 0.013° 2Θ with a step-counting time of 0.5 s was used in a scanning range from 3° to 80° 2Θ. Semi-quantitative mineral content was calculated based on relative intensity values using the software package DIFRAC.EVA V2 (Bruker, USA). Absolute reproducibility variation was < 1% (n = 4).

### M6. DNA extraction, Illumina library preparation and sequencing

Metagenomic DNA was extracted from 10 g of soil as described previously (4). Briefly, the soil was mixed for 30 minutes in 40 mL cell extraction buffer (1% PEG 8000 ; 1M NaCl, pH 9,2) (5). The supernatant was ultra-centrifuged 2 h at 44,000 x g at 4 °C and DNA was extracted from the pellet using a bead-beating and phenol/chloroform/isoamylalcohol based protocol (6). DNA was resuspended in 30 µL of DEPC treated water. Two extractions were performed per sample and the resulting DNA was combined. DNA concentration was measured using the Qubit 1x dsDNA HS Assay Kit (Thermo Fisher Scientific) and Qubit 4 Fluorometer (Thermo Fisher Scientific). 10 mL of the cell extraction buffer was used as a negative control for DNA extraction.

Approximately 5-15 ng of DNA were shared with a E220 Focused-ultrasonicator (Covaris® Inc., MA, USA), targeting 300-400 fragment size, and used to prepare the metagenomic libraries. The libraries were constructed using the NEBNEXT® ultra II DNA library prep kit for Illumina and the NEBNext® primer set 1 (Dual index, New England BioLabs, UK) with three modifications. 1) the primer adapters were diluted 1:50 v/v, 2) the primers were diluted 1:2 v/v and 3) a second cleaning step was performed after PCR amplification. Purification and size selection were conducted using magnetic beads Agencourt® AMPure® XP (Beckman-Coulter, MA, USA). Inserts between 400 and 500?bp were kept and their quality evaluated using a Fragment Analyzer™ (Advanced Analytical, IA, USA). Library concentration was measured with the Qubit 1x dsDNA HS Assay Kit and Qubit 4 Fluorometer. The metagenomic libraries were sequenced on an Illumina HiSeq 2500 (Illumina, CA, USA) using the HiSeq Rapid SBS Kit v2 (500 cycles, Illumina, CA, USA) and loading 12 pM including 1% v/v PhiX.

### M7. Metagenome assembly, binning and annotation

HiSeq reads were quality filtered using BBduk (https://sourceforge.net/projects/bbmap/) and sickle (https://github.com/najoshi/sickle). metaSPADES 3.13 (7) was used to assemble the reads and the resulting scaffolds were filtered for length >= 1000 bp for gene prediction using Prodigal (8) in meta mode and annotation using Diamond version 0.9.9 (9) against the UniRef100 database (10) with e-value cut-off of 1E-5. Scaffold coverages were calculated by mapping reads using Bowtie2 in sensitive mode (11). Genomes were binned using abawaca (github.com/CK7/abawaca), ESOM (12) and MaxBin2 (13), and the resulting bins were aggregated using DAS Tool (14). Each genomic bin was manually curated using coverage, gene-based taxonomy and GC content information for each scaffold. ra2 (15) was used to fix assembly errors in all binned scaffolds. CheckM (16) was used to estimate the quality of the bins and only high quality bins with completeness >75% and contamination <15% were considered for further analysis. For all high quality genomes, GTDB-Tk classify_wf (17) was used for a broad taxonomic classification and *in situ* genome replication measures (iRep) (18) were calculated using --mm 3 flag after mapping the reads to scaffolds with Bowtie2 (19). Further functional and metabolic capacities of high quality genomes and metagenomes were determined using METABOLIC (20). METABOLIC output was further expanded upon using hidden-markov-model (HMM) search results for the archaeal amoA protein (HMMER v3.2 (http://hmmer.org/), -Z 47079205 -E 1000) and other genes previously annotated using the UniREF100 database (10). Relative abundances of key metabolic genes were calculated by identifying scaffolds carrying the gene in question, summing up their coverages and finally normalizing the summed coverage with the sequencing depth of each respective sample.

## Supplementary Results and Discussion

### Extended pangenome analysis

Notably, none of the eight Thaumarchaeota genomes contained CRISPR arrays with more than one spacer and only ABT-LB2 contained a putative *cas* gene. Similarly no Cas gene was found in *N. franklandus, N. oleophilus* and *N. exaquare*, and one Cas gene was found in N. arcticus. Only *N. exaquare* contained an evidence level 4 CRISPR array with 5 spacers (**Table S11**). The lack of CRISPR-Cas systems in these environments could be due to the lower presence of *Thaumarchaeota* targeting viruses in these environments and/or be coupled with slow growth rates rendering the CRISPR-Cas system immune response ineffective (21).

Electron transfer flavoprotein fixABCX genes were found only amongst the ABT genomes and not in any of *Ca. Nitrosocosmici* (**Table S11**). These genes are reported to be involved in the electron bifurcation in diazotrophs (22), but are also found in many non-diazotrophic *Archaea* (23), where their function is yet to be determined.

No S-layer protein slp1 was found in any of the ABT nor other *Ca. Nitrosocosmici*. The presence of Hexuronic acid methyltransferase AglP, which is involved in the pathway of S-layer biogenesis, suggests that there may exist an alternative pathway for S-layer production. This could provide additional protection against harsh desert environments for the ABT genomes (24).

## Supplementary Figures and Tables

**Table S1:**
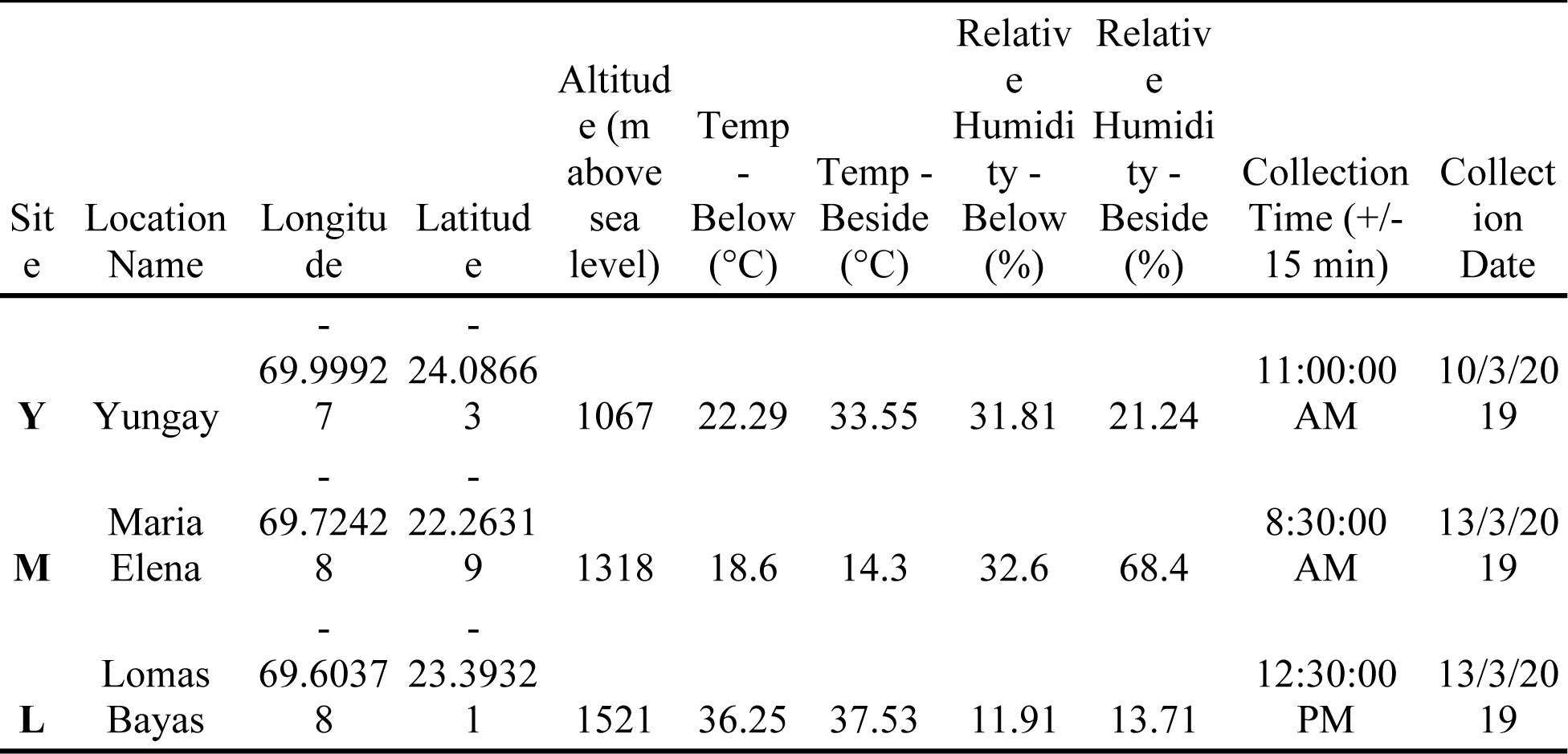
Sampling information, temperature and relative humidity below and beside boulders were measured using OBO U23 pro temperature/relative humidity data logger at the time of sampling.

**Table S2.**
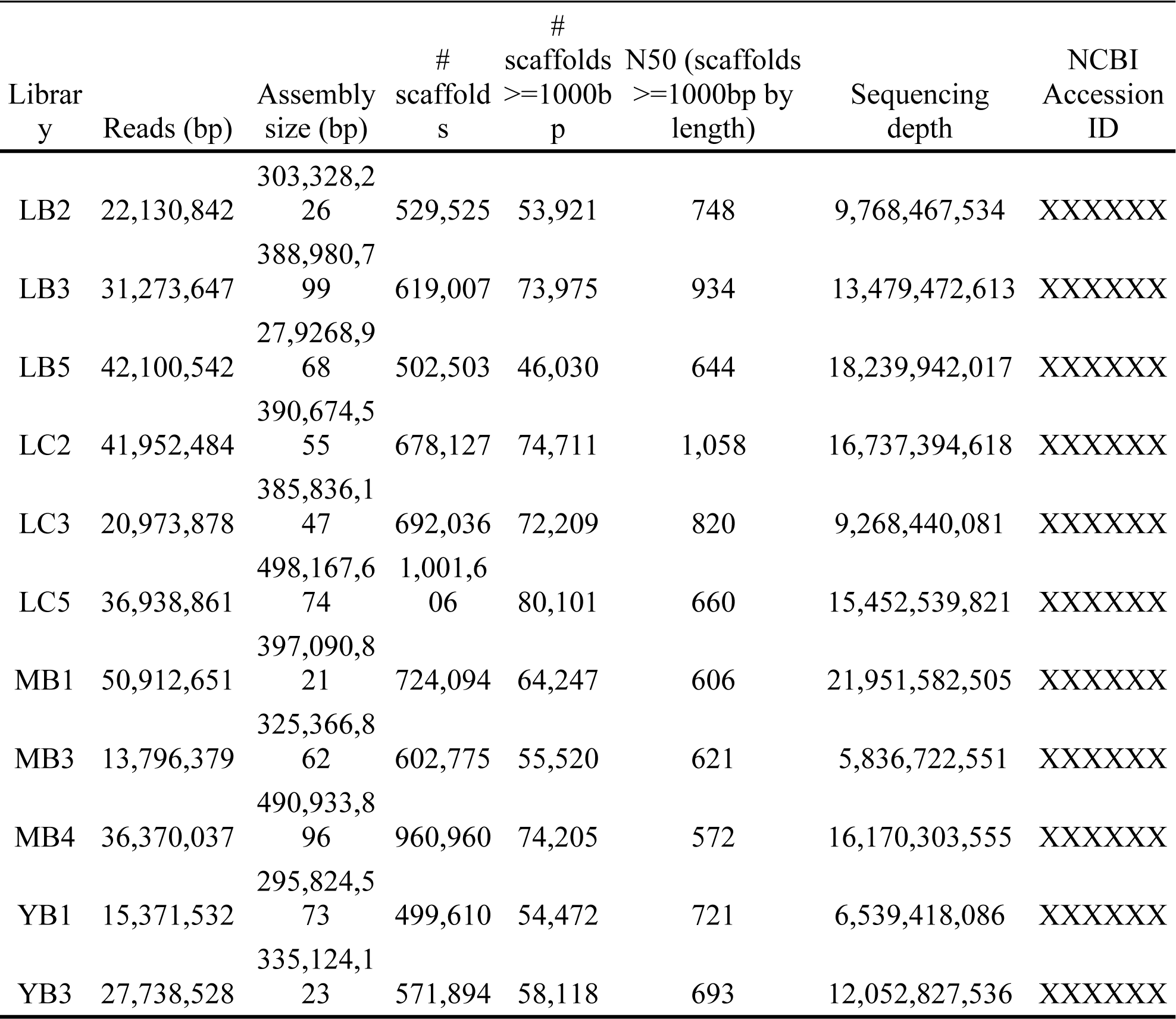
Metagenome library information. DNA extracts from MB6, MC3, YB2 and YB5 contained measurable DNA (see **Figure S1**) however, failed in library preparation.

**Figure S1.**
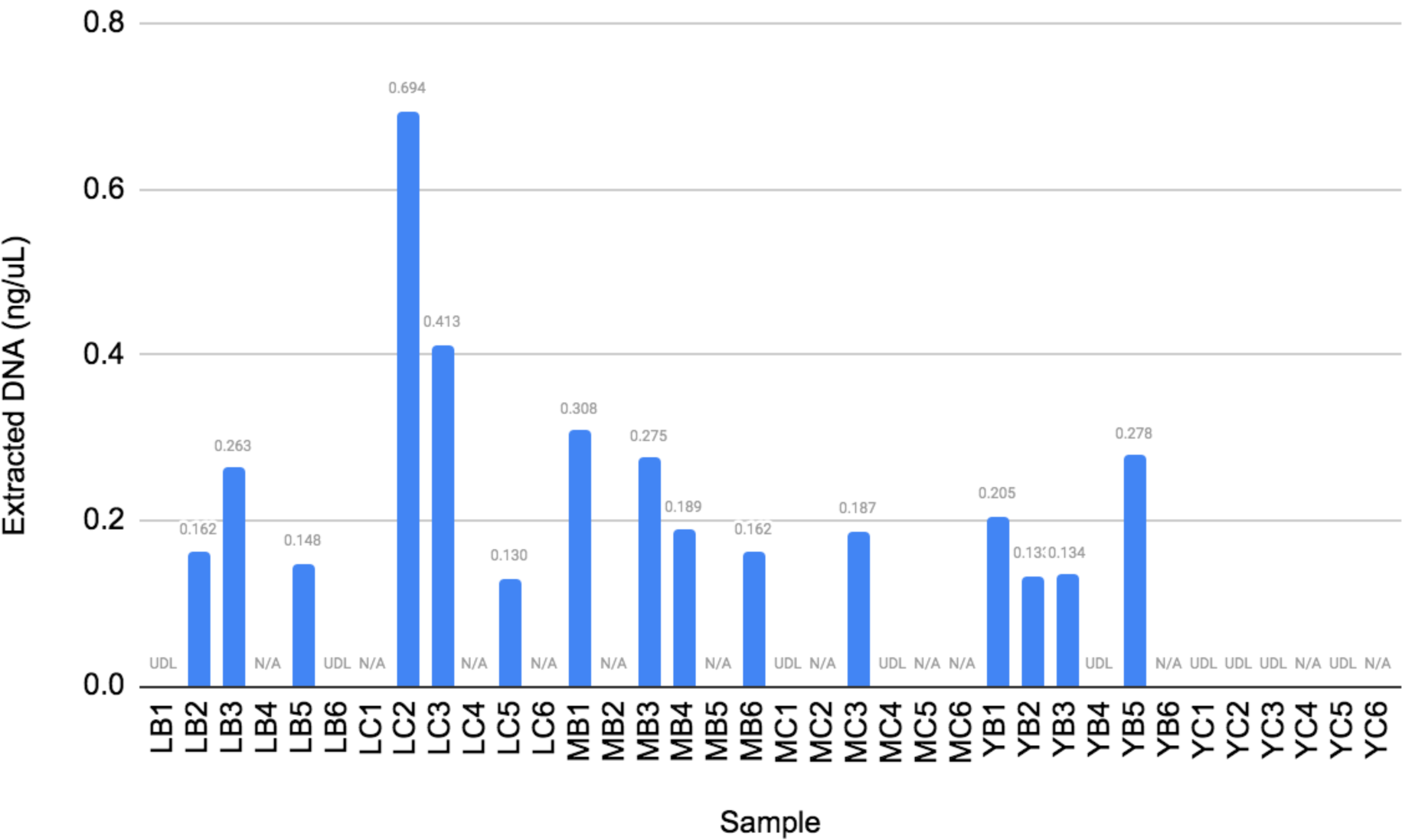
DNA extraction results. UDL (under detection limit) indicates extraction was attempted but resulted in DNA amount below detection limit (0.01 ng/uL) and N/A indicates no extraction was attempted.

**Figure S2.**
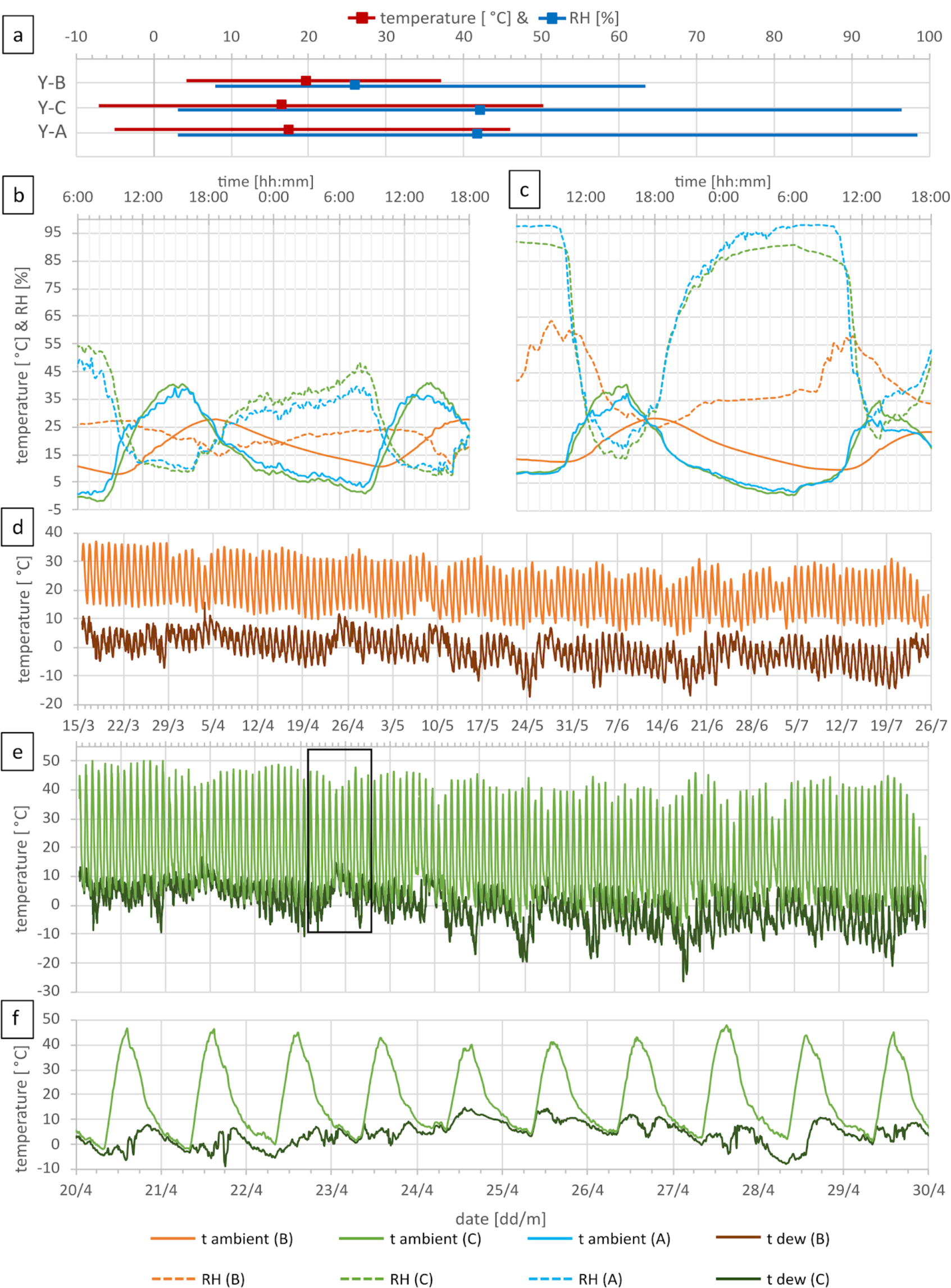
Extended field measurements of temperature and relative humidity (RH) for Y-below boulder (B), control (C) and in 1 m above ground (A). a) Mean temperature and RH (square) and range (bar). b) Temperature and RH during a dry diurnal cycle; c) Temperature and RH during a moist diurnal cycle. d) Ambient temperature and calculated dew point temperature below boulder (B) during the full 130 days of recording. e) Ambient temperature and calculated dew point temperature beside boulder (control) during the full 130 days of recording, black rectangle is zoomed in panel f).

**Figure S3.**
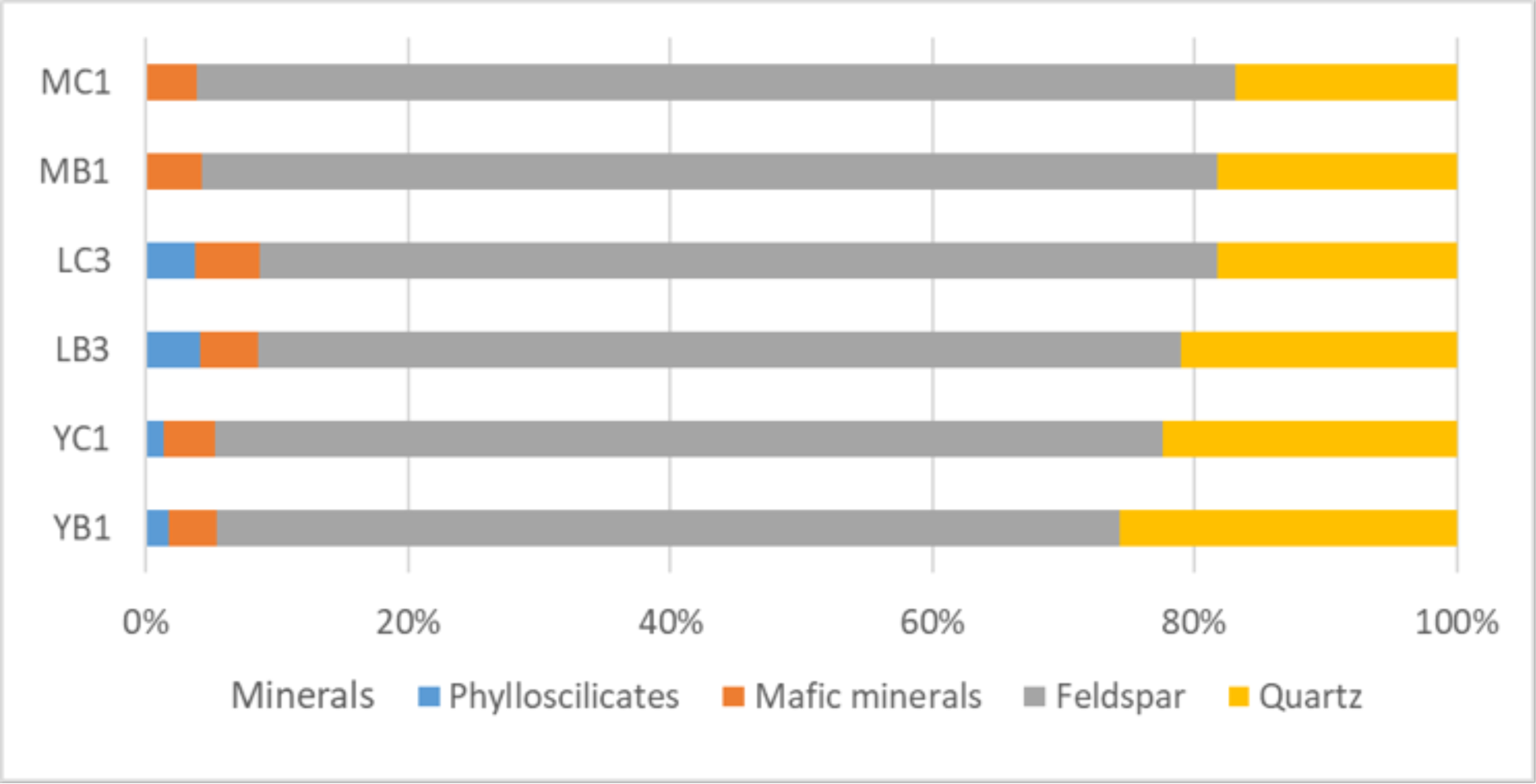
Mineral composition determined using XRD.

**Figure S4.**
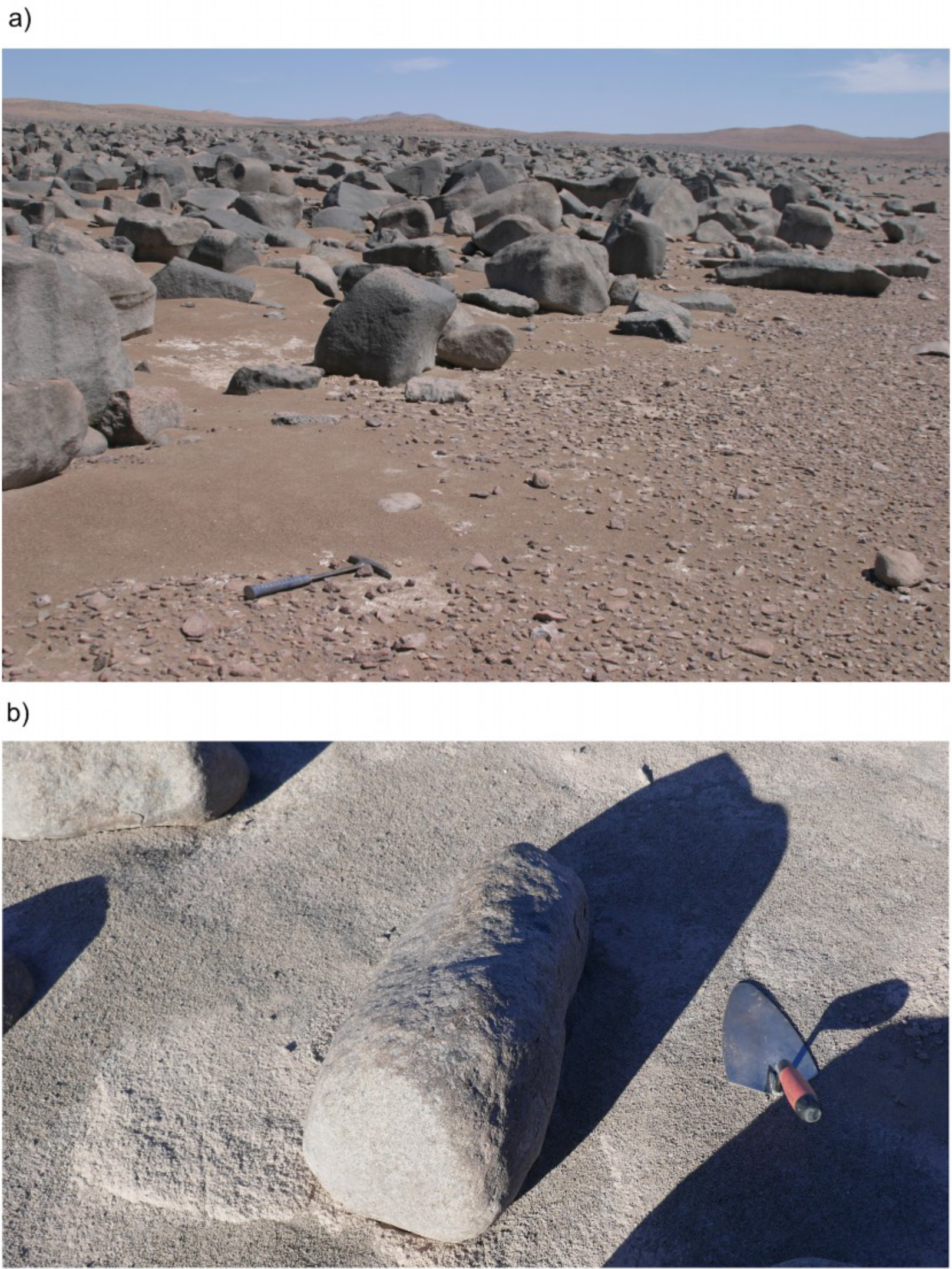
Atacama Boulder Fields. a) Yungay Valley boulder field. b) An example of the boulders chosen for sampling.

**Figure S5.**
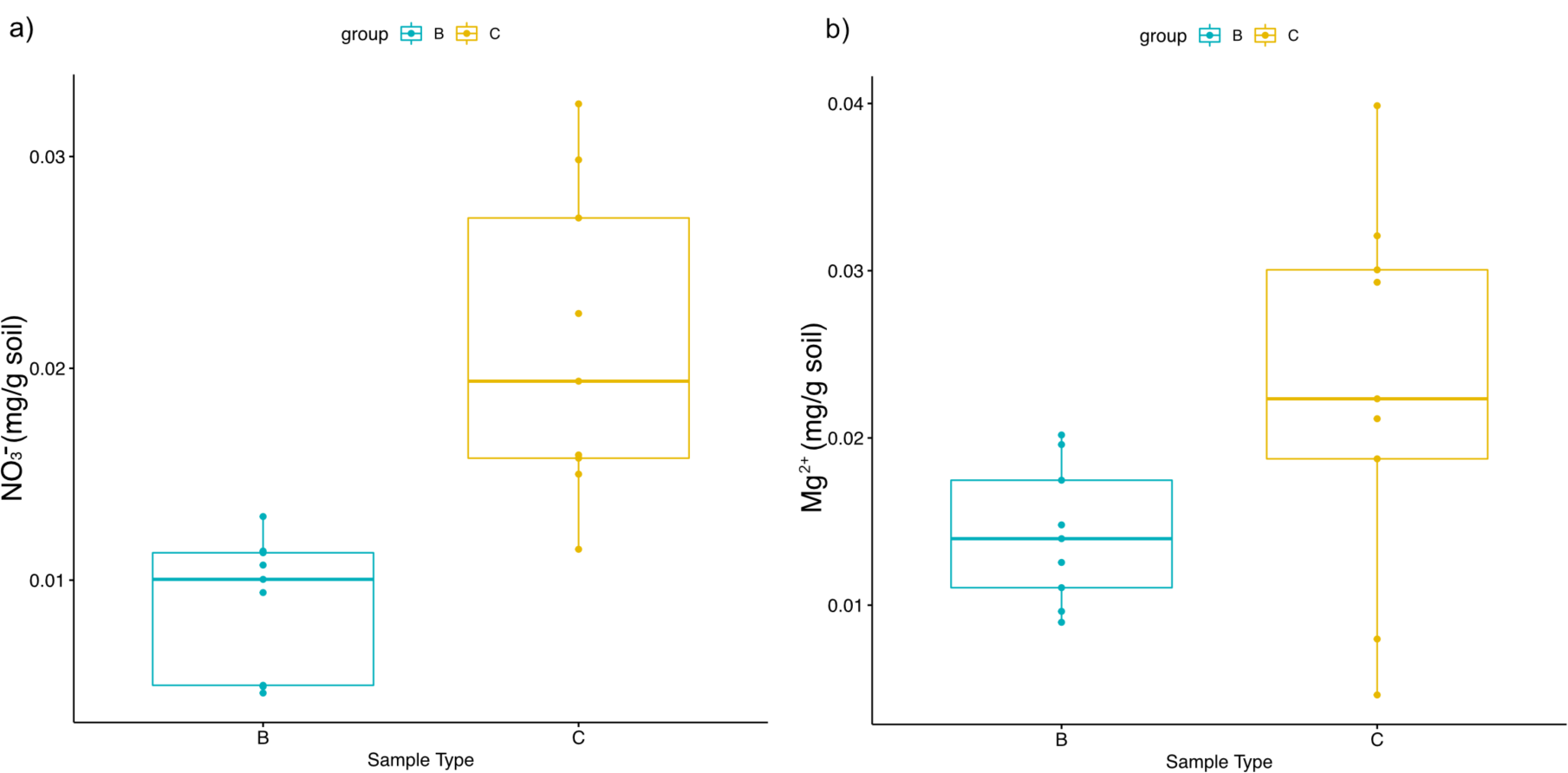
Comparison of a) nitrate and b) magnesium ion concentrations between B and C sample types. Plots were visualized using “ggpubr” package in R.

**Figure S6.**
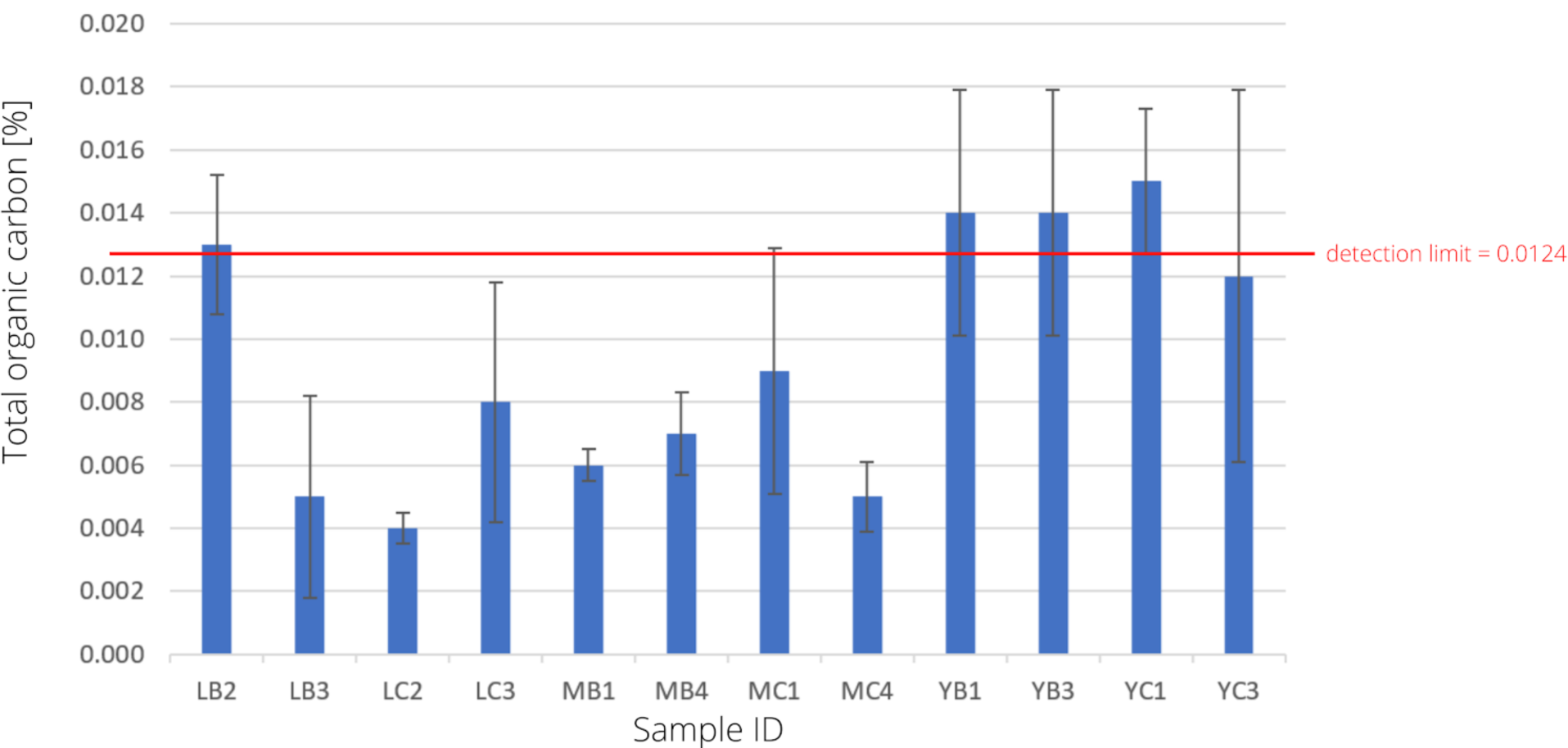
Total Organic Carbon (TOC) content [wt%] per sample. All values were below the limit of quantitation value 0.02962 wt% and very close to or below the limit of detection 0.0124 wt%.

**Figure S7.**
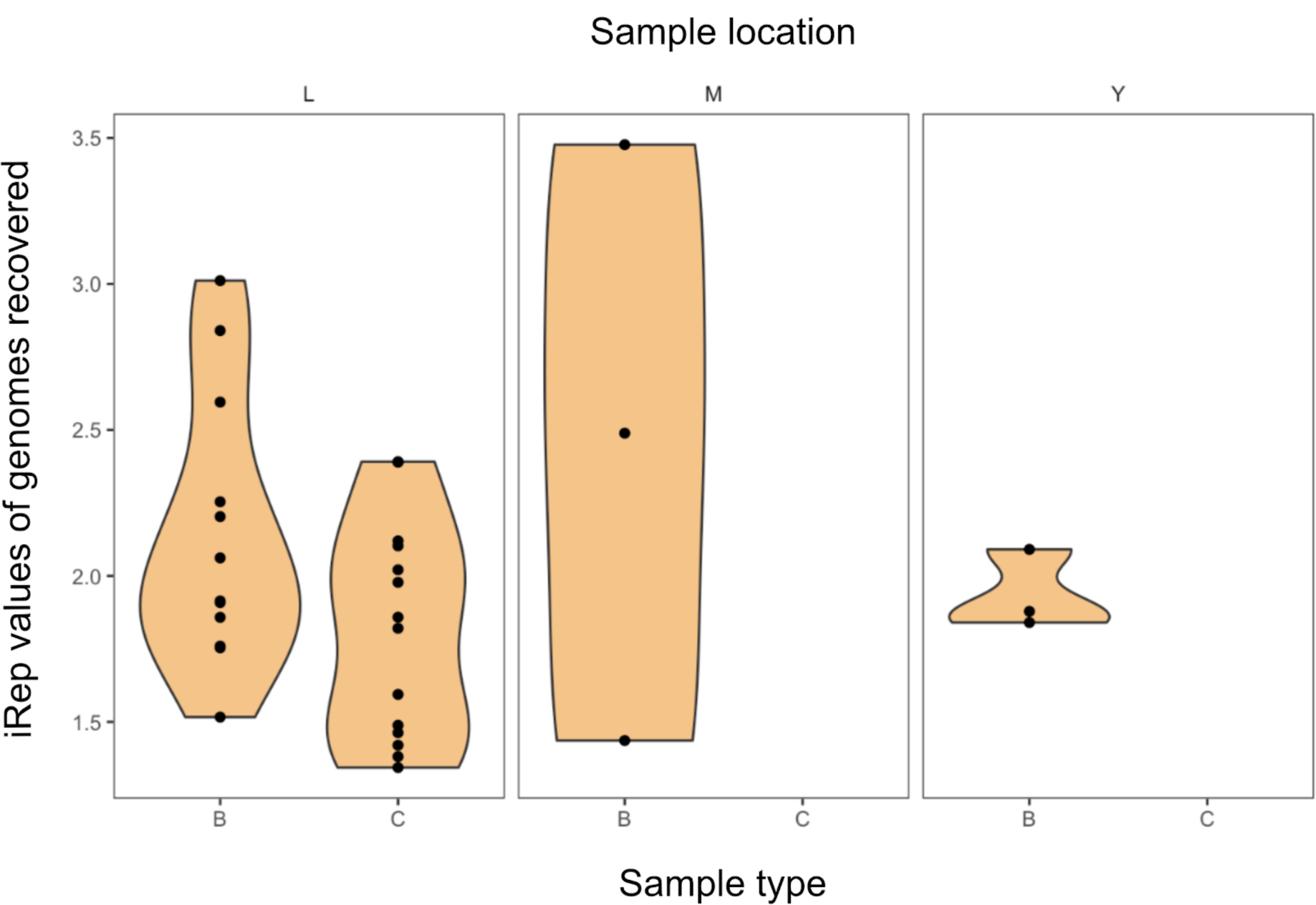
calculated iRep values of genomes of sample type (B and C) and sample sites (L, M and Y). Plot was visualized using ggplot2 (25)

**Figure S8.**
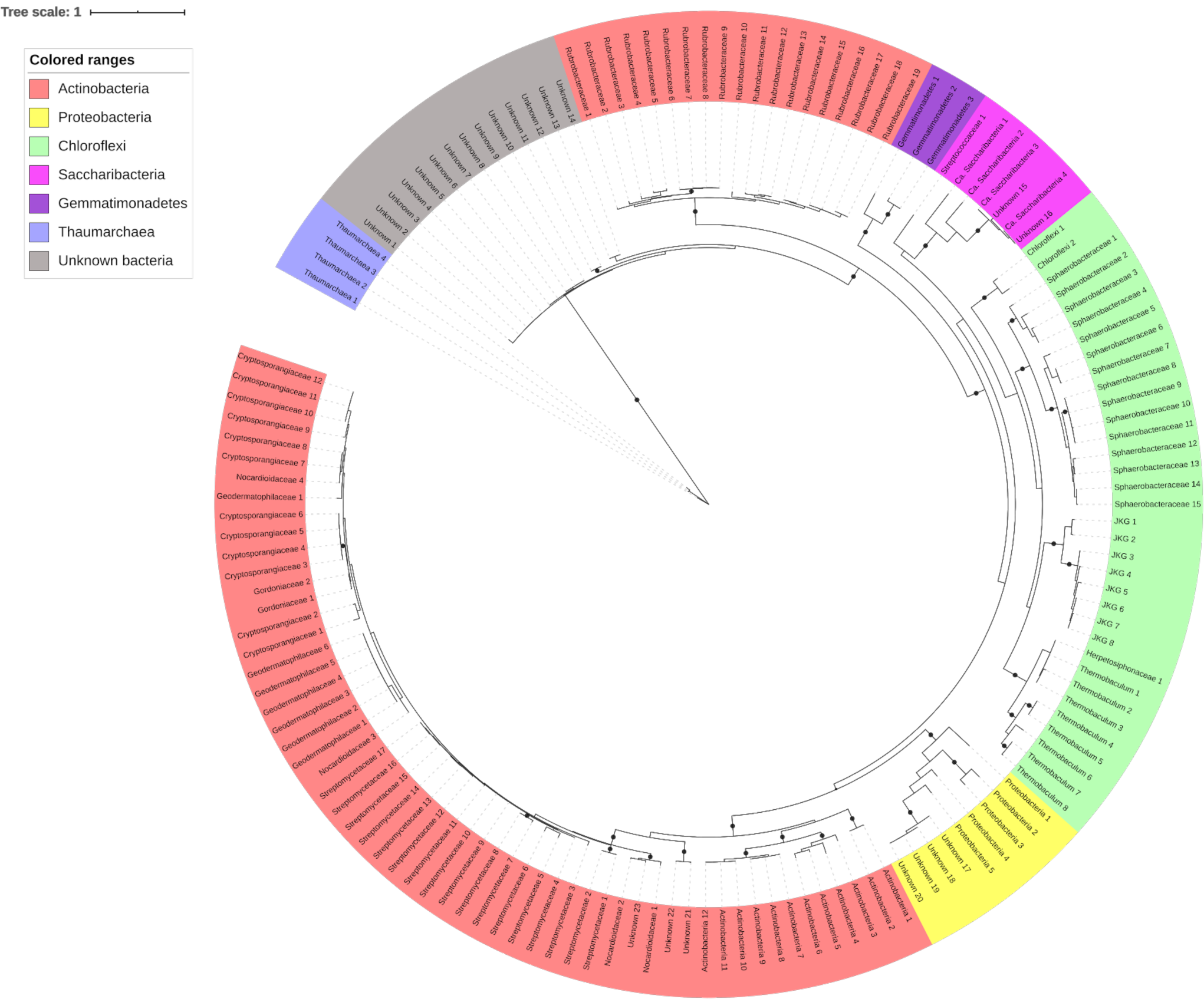
Full phylogenetic tree of all recovered rpS3 gene clusters. Color ranges refer to phyla level classification, leaf labels refer to taxonomic resolution down to family level based on BLAST (26) results against UniRef100 (10). Strongly supported branches as described in the Methods section are indicated with black dots.

**Figure S9.**
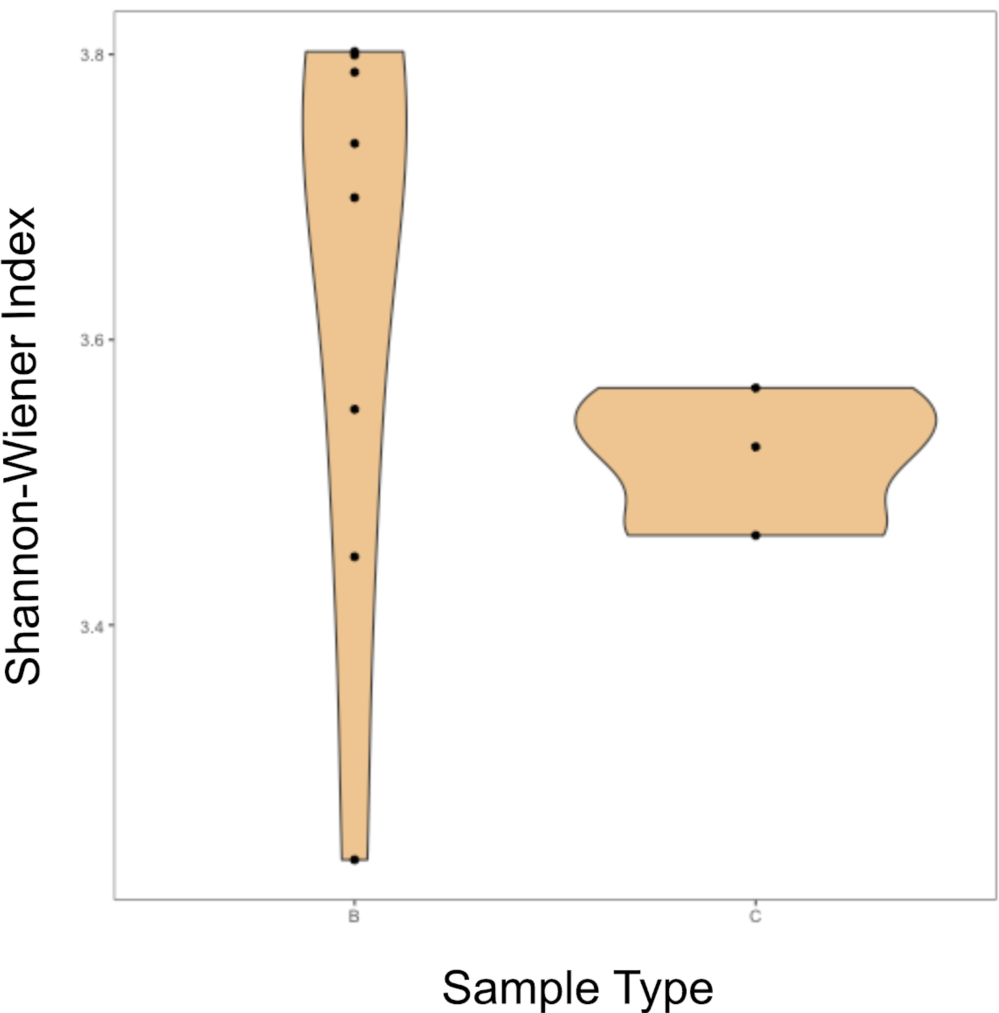
Shannon-Wiener index of each metagenome. Indices were calculated based on normalized ribosomal protein S3 (rpS3) abundances. Plot was visualized using ggplot2 (25).

**Figure S10.**
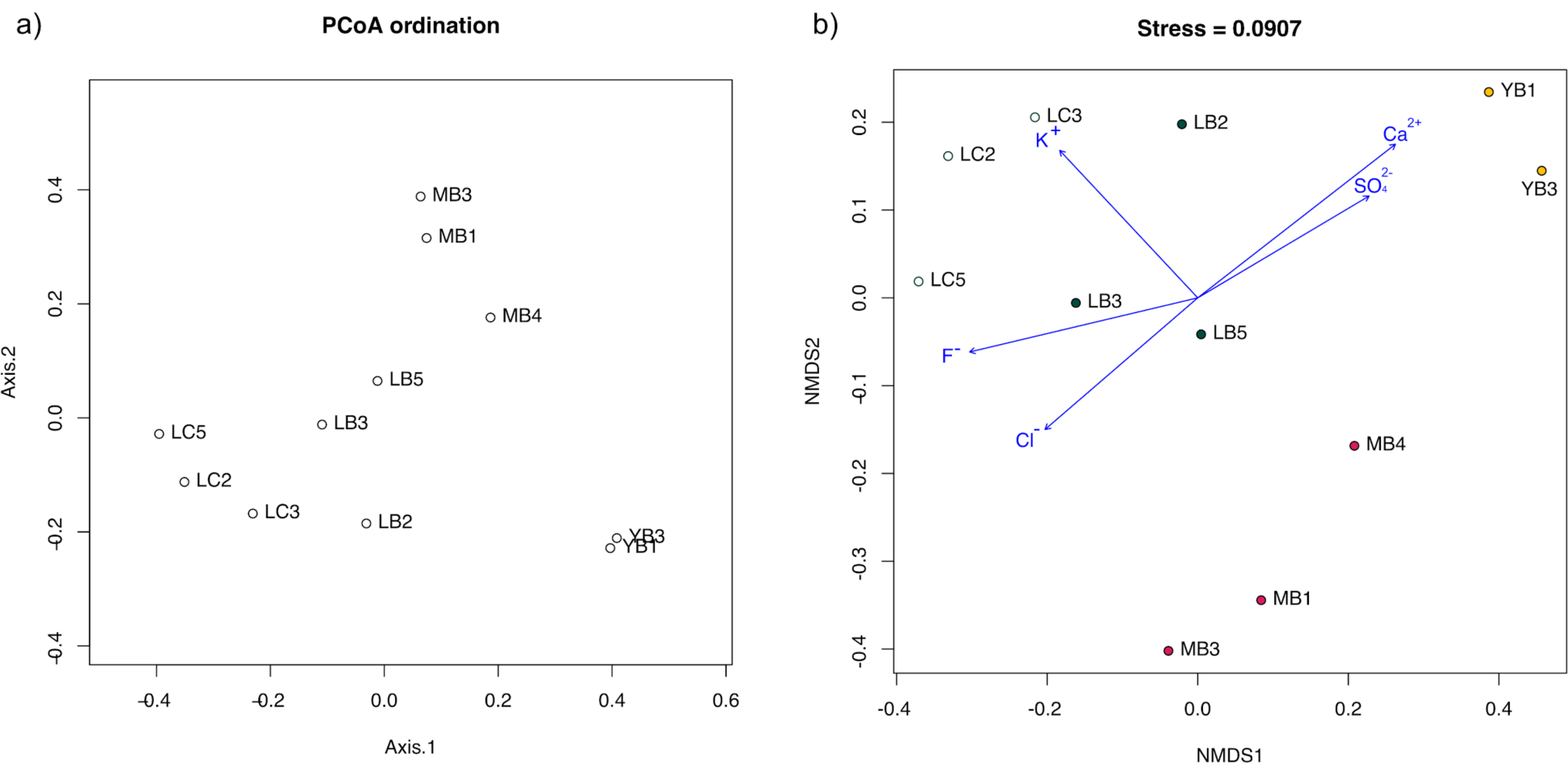
a) PCoA and b) NMDS ordination plots of metagenomes based on rpS3 abundances. Bray-Curtis distance matrix of normalized abundances of rpS3 taxa across all metagenomes were calculated and used as input for both figures. For NMDS, ion concentration meta data was added and the vectors were fitted with the ordination. Blue arrows represent fitted ion species with a p-value less than 0.1. Both figures were generated using R.

**Figure S11.**
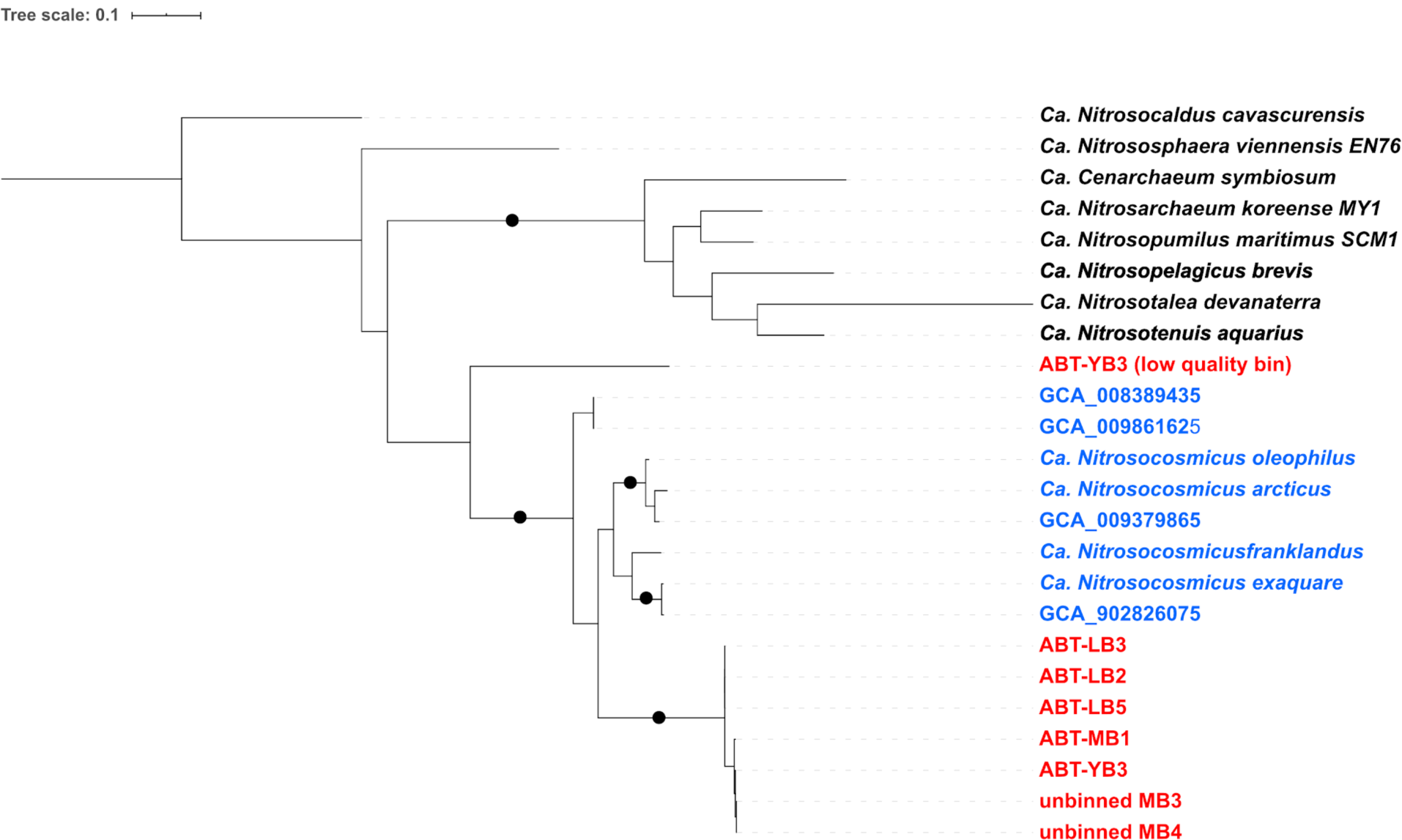
Tree of all *amoA* sequences from this study (red), *amoA*s from *Ca. Nitrosocosmicus* (Blue) and other representative Thaumarchaea sequences (Black). Strongly supported branches as described in the M&M section are indicated with black dots.

**Figure S12.**
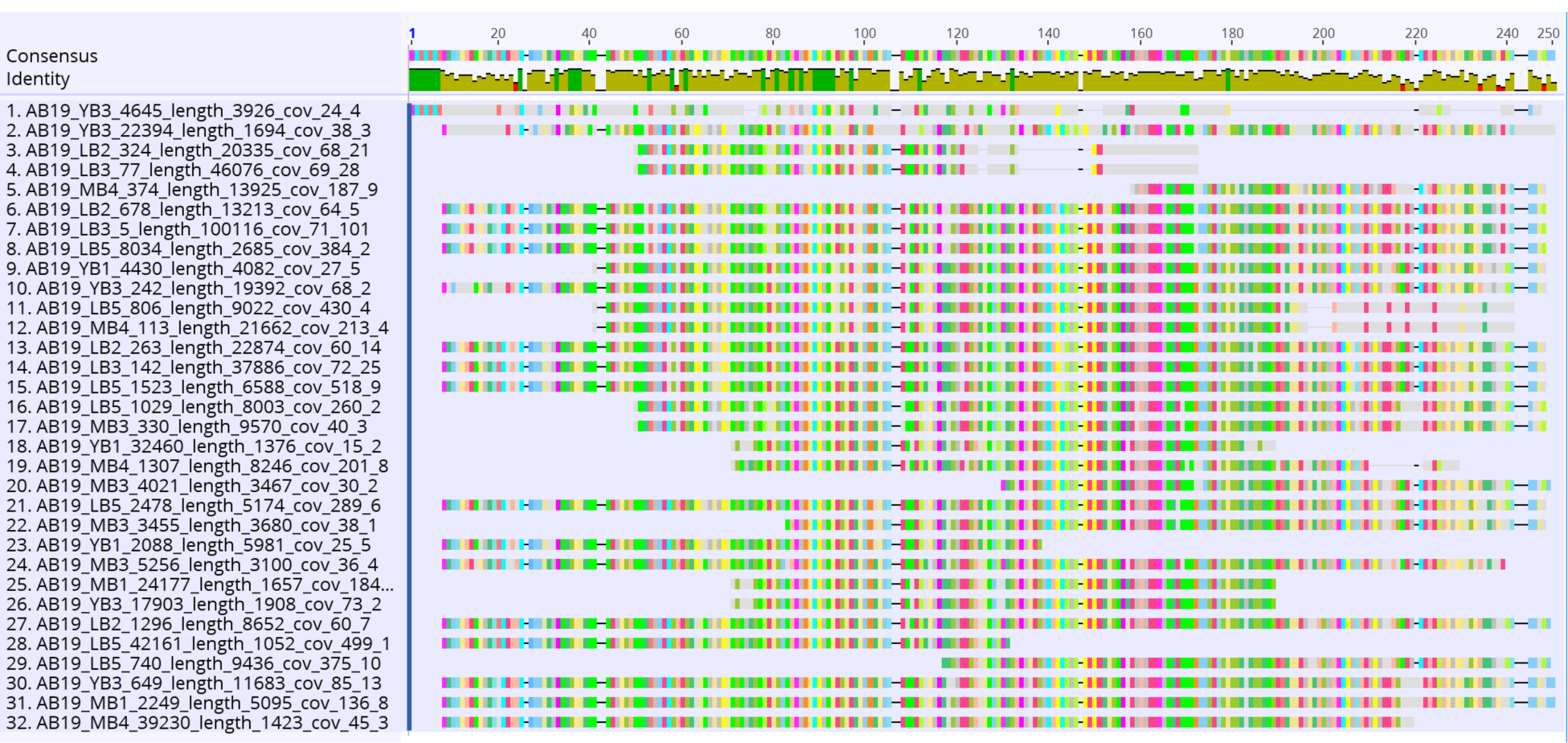
Alignments of 32 aquaporins recovered across all samples. Visualization using the Geneious software. Sequences 1-4 show a high level of sequence divergence, while the rest show truncation at both ends despite most being located mid scaffold.

**Figure S13.**
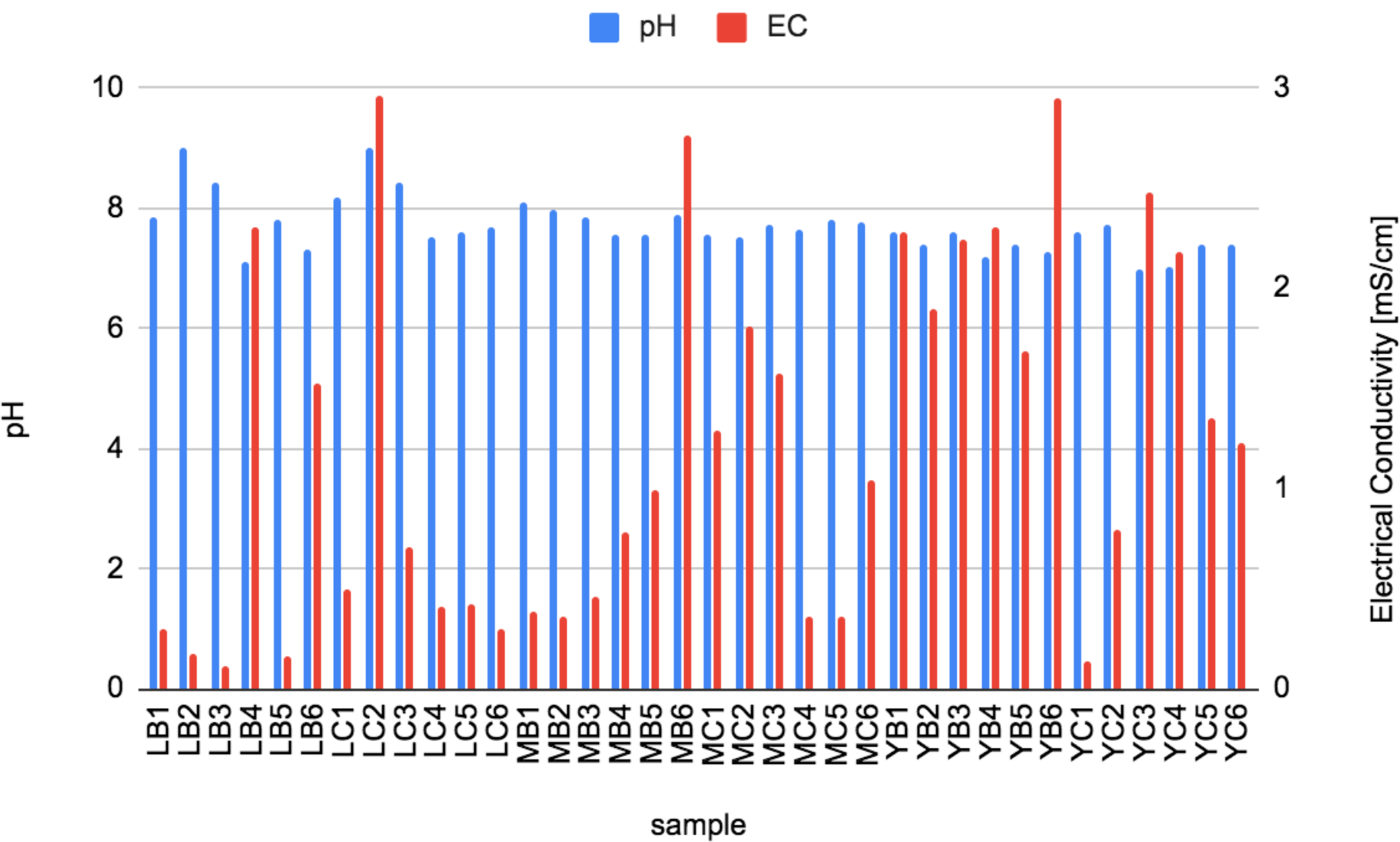
pH and electrical conductivity (EC) of each sample.

**Tables S3 - S14 are available as a separate Excel file:**

**Table S3. Statistics and meta-data of the reference genomes used for comparative genomics**.

**Table S4. Genome statistics of high quality genomes**. High quality genomes were determined using CheckM completeness > 75 % and contamination < 15 %. CheckM(16) output (Completeness, contamination, GC std, # ambiguous bases, Genome size, Longest Contig, N50 (scaffolds), Mean scaffold length, # contigs, # scaffolds, # predicted genes, Longestscaffold, GC, N50 (contigs), Coding density, Mean contig length) are accompanied by iRep value (18)) for genomes whose iRep values could be calculated, rpS3 taxa based on BLAST(26) results against UniRef100 database (10), gtdb-tk classification (17) using “classify_wf”.

**Table S5. Normalized abundances of all rpS3 taxa across all samples**. Column 1 corresponds to the scaffold containing the centroid of each rpS3 cluster, which was then used to calculate the normalized abundance of each rpS3 cluster (taxa) across all samples.

**Table S6. ANOVA p-values of all rpS3 taxa**. For all 147 rpS3 taxa ANOVA tests were performed between Boulder and Control sample groups (B x C), YB and MB samples (YB x MB), YB and LB samples (YB x LB), and LB and MB samples (LB x MB)

**Table S7. METABOLIC output of metagenomes**. Presence, count and gene ID of all the predicted metabolic genes found across the metagenome.

**Table S8. METABOLIC output of all high quality genomes**. Presence, count and gene ID of all the predicted metabolic genes found across the metagenome.

**Table S9. Ammonification genes**.

**Table S10. Orthologous protein clusters with GO** (27,28) **and Swiss-Prot**(29) **annotation**. Clusters were determined using Orthovenn2(30). In orange are clusters with putative functions associated with stress response and in yellow are clusters with putative functions associated with nitrogen metabolism.

**Table S11. List of amoABCX genes, 4HB/3HP pathway genes, TCA cycle, gluconeogenesis, pentose phosphate pathway and other notable genes for each ABT genomes**.

**Table S12. Singletons of LB3, MB4, YB3 genomes**. Singletons were identified using Orthovenn2(30) and annotated by BLASTing (26,30) against UniRef100 database (10)

**Table S13. List of NCBI genomes used for phylogenomic tree construction**. NCBI genomes classified as Thaumarchaeota on 30th May 2020, filtered using CheckM completeness >50 % and contamination < 5%.

**Table S14. Ion chromatography raw data in mg/g soil**.

## Additional File

**Additional File 1**: Newick treefile of *Ca. Nitrosodesertus* and NCBI genomes annotated as *Thaumarchaeaota*

## Notes

### Competing Interest Statement

The authors have declared no competing interest.

